# Impaired photoprotection in *Phaeodactylum tricornutum* KEA3 mutants reveals the proton regulatory circuit of diatoms light acclimation

**DOI:** 10.1101/2021.09.06.459119

**Authors:** Claire Seydoux, Mattia Storti, Vasco Giovagnetti, Anna Matuszyńska, Erika Guglielmino, Xue Zhao, Cécile Giustini, Yufang Pan, Jhoanell Angulo, Alexander V. Ruban, Hanhua Hu, Benjamin Bailleul, Florence Courtois, Guillaume Allorent, Giovanni Finazzi

## Abstract

Diatoms are amongst the most successful clades of oceanic phytoplankton, significantly contributing to photosynthesis on Earth. Their ecological success likely stems from their ability to acclimate to changing environmental conditions, including e.g. variable light intensity. Diatoms are outstanding at dissipating light energy exceeding the maximum photosynthetic electron transfer (PET) capacity of via Non Photochemical Quenching (NPQ). While the molecular effectors of this process, as well as the role of the Proton Motive Force (PMF) in its regulation are known, the putative regulators of the PET/PMF relationship in diatoms remain unidentified. Here, we demonstrate that the H^+^/K^+^ antiporter KEA3 is the main regulator of the coupling between PMF and PET in the model diatom *Phaeodactylum tricornutum*. By controlling the PMF, it modulates NPQ responses at the onset of illumination, during transients and in steady state conditions. Under intermittent light KEA3 absence results in reduced fitness. Using a parsimonious model including only two components, KEA3 and the diadinoxanthin de-epoxidase, we can describe most of the feedback loops observed between PET and NPQ. This two-components regulatory system allows for efficient responses to fast (minutes) or slow (e.g. diel) changes in light environment, thanks to the presence of a regulatory Ca^2+^-binding domain in KEA3 that controls its activity. This circuit is likely finely tuned by the NPQ effector proteins LHCX, providing diatoms with the required flexibility to thrive in different ocean provinces.

**One sentence summary:** The author(s) responsible for distribution of materials integral to the findings presented in this article in accordance with the policy described in the Instructions for Authors (https://academic.oup.com/plcell/pages/General-Instructions) is Giovanni Finazzi.

## Introduction

Diatoms are key ecological players and very efficient CO_2_ sinks in contemporary oceans (Tréguer et al., 2018; Falciatore et al., 2020). They thrive in diverse environmental niches (Malviya et al., 2016), proliferate on ice (Thomas and Dieckmann, 2002) and generally dominate plankton communities in upwelling turbulent environments (Margalef, 1978; Tréguer et al., 2018; Falciatore et al., 2020). A likely key factor for diatom ecological success in challenging environments is their photosynthetic flexibility. Photosynthetic electron flow is optimized by the peculiar 3D organization of photosynthetic complexes within the thylakoid membranes, which are organized as parallel interconnected layers (Flori et al., 2017). Light energy is collected by a specific peculiar light-harvesting apparatus (Büchel, 2020), in which specific pigments (i.e. chlorophyll *a* and *c*, β-carotene, fucoxanthin, diadinoxanthin -DD, diatoxanthin -DT) are embedded in antenna complexes, which either absorb sunlight (LHCF and LHCR families) or mediate the photoprotective NPQ response (LHCX family, (Büchel, 2020; Falciatore et al., 2020)). Diatoms display very high NPQ levels (Lavaud et al., 2002c; Ruban et al., 2004), triggered by the reversible enzymatic conversion of DD into DT through the xanthophyll cycle (XC, (Stransky and Hager, 1970; Jakob et al., 2001; Blommaert et al., 2021)). Accumulation of violaxanthin-antheraxanthin-zeaxanthin (the XC pigments of plants) is also observed under specific conditions (Lohr and Wilhelm, 1999). In diatoms, NPQ responses are modulated by the LHCXs proteins (Bailleul et al., 2010; Lepetit et al., 2017; Buck et al., 2019), which are NPQ effectors like the PSBS and LHCSRs proteins in *Viridiplantae* (Niyogi and Truong, 2013; Giovagnetti and Ruban, 2018; Büchel, 2020; Falciatore et al., 2020).

In *Viridiplantae*, the Proton Motive Force (PMF) sets the NPQ response (Wraight and Crofts, 1970), activating the XC enzymes and inducing the protonation of specific amino acids on PSBS and LHCSRs during ΔpH formation (Li et al., 2004; Zaks et al., 2012; Niyogi and Truong, 2013; Matuszyńska et al., 2016; Ballottari et al., 2016; Davis et al., 2017). In diatoms, the role of the PMF on NPQ regulation is less understood: the XC is pH-sensitive (Jakob et al., 2001; Blommaert et al., 2021), while although a pH-related modulation of the function of LHCX proteins cannot be ruled out, it is less likely to occur (Falciatore et al., 2020; Shinkle et al., 2010). This difference could reflect diverse features of LHCX vs. PSBS/LHCSR proteins, with less numerous protonable residues facing the thylakoid lumen in the former (Taddei et al., 2016). It has been proposed that the mechanism of DD-dependent NPQ does not depend on the presence of the ΔpH and that DT is able to exert its full quenching capacity in the absence of a proton gradient (Goss et al., 2006). This proposal was criticized by Lavaud and Kroth (Lavaud and Kroth, 2006), who proposed the existence of a pH control beyond the XC. Given the crucial role of PMF on diatom’s NPQ, here we re-examined the coupling between photosynthetic activity and the PMF, using photophysiology, genetic and modelling approaches in the model species *Phaeodactylum tricornutum*. We found that the luminal pH, which regulates NPQ through the pH dependence of the DD de-epoxidase, is in turn modulated by the H^+^/K^+^ antiporter KEA via a parsimonious two-step regulatory circuit. An EF hand motif, which we could identify in the *P. tricornutum* KEA3 but not in the *Viridiplantae* orthologs, was shown to control its activity, providing a possible link between intracellular Ca^2+^ and responses to fast (minutes) or slow (e.g. diel) changes in light environment.

## Results

### NPQ is pH dependent in diatoms

We investigated the pH dependency of diatoms NPQ in *P. tricornutum* using a previously established protocol to equilibrate the pH across algal internal compartments (Finazzi and Rappaport, 1998; Tian et al., 2019). Cells were exposed to a progressive acidification of the external medium pH in the presence of a permeant buffer (acetate). pH equilibration was facilitated using the H^+^/K^+^ exchanger nigericin. Lowering the pH induced a significant fluorescence quenching (Figure 1A), similar to the one observed in untreated cells upon exposure to high light (HL). The pH-induced quenching in the dark was accompanied by the conversion of DD into DT (Figure 1B, red line) to a similar extent of that observed during light-induced NPQ in non-permeabilized cells (Figure 1B, black line). Initial fluorescence levels were recovered upon reversing the external pH to its initial neutral level with KOH (Figure 1A, B). We observed the same pH dependency of NPQ in two different ecotypes of *P. tricornutum*: Pt1 and Pt4 (Supplementary Figure S3), which display different levels of NPQ.

**Figure 1.**
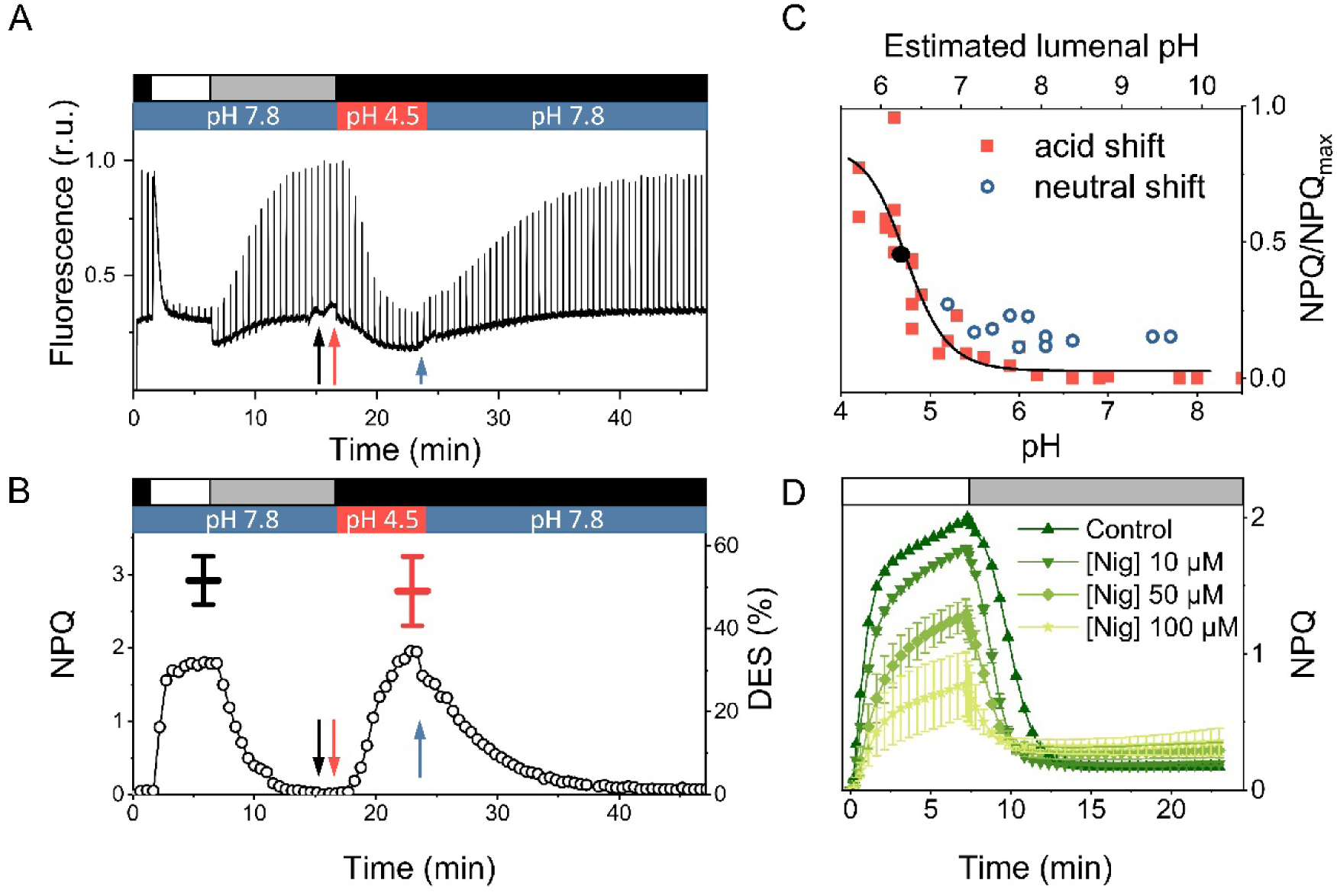
NPQ in *Phaeodactylum tricornutum* is pH and ΔpH dependent. (**A**) Fluorescence transients in *P. tricornutum* WT cells exposed to HL (1200 µmol photons m^-2^ s^-1^, white box), LL (25 µmol photons m^-2^ s^-1^, grey box) and dark (black box) at different pH values. Arrows indicate addition of nigericin (20 µM, black), acetic acid (5 mM, red) and KOH (2,5 mM, blue). (**B**) NPQ kinetics (circles) calculated from the fluorescence traces shown in panel (A). Black line: De-epoxidation State (DES) in HL. Red line: DES at pH 4.5 (in the dark). N = 4. Mean ± SD. (**C**) pH dependency of NPQ in the presence of 20 µM nigericin during an acidic (red squares) or a neutral (blue circles) shift. Black dot: apparent pKa of the NPQ vs pH relationship. The acid shift was done from an initial external pH of 8 to the indicated pH values. The pH neutral shift was done from an initial external pH of 4.5 to the indicated pH values. Data represent different experiments performed with 6 biological samples. (**D**) Nigericin sensitivity of NPQ in *P. tricornutum* cells. Different concentrations of nigericin were added before measuring NPQ induction in HL (1200 µmol photons m^-2^ s^-1^, white box) and relaxation in LL (25 µmol photons m^-2^ s^-1^, grey box). N = 3. Mean ± SD.

The absolute pH dependency of quenching in *P. tricornutum* (apparent pKa of 4.7, Figure 1C) was shifted towards more acidic values than in green algae (apparent pKa of 6.2 in *Chlamydomonas reinhardtii*, (Tian et al., 2019)). Thus, quenching could reflect photosystem II (PSII) photodamage, which is expected at such low lumen pH values (Krieger and Weis, 1993; Spetea et al., 1997; Kramer et al., 2003). However, we found a very similar Stern-Volmer (S-V) relationship between NPQ and PSII photochemical quantum yield (Fv’/Fm’) during a HL to LL relaxation and the acid to neutral pH transition (Supplementary Figure S1). This suggests that PSII maintained the same photochemical capacity in HL and during the pH shift. Thus, the different pH dependence could reflect, at least in part, an incomplete pH equilibration between the lumen and the external medium during the experiment. Indeed, a calibration of the luminal pH during the pH shift using an independent probe (the pH dependence of cytochrome *b_6_f* turnover, Supplementary Figure S2) pointed to pH luminal values higher than the ones imposed to the cell medium. Despite this possible inaccuracy in assessing the absolute NPQ/pH relationship in permeabilized cells, the effect of acidification on fluorescence quenching demonstrates the role of the pH component of the PMF on diatoms NPQ. This notion was corroborated by testing how *P. tricornutum* NPQ kinetics is affected by adding two protonophores, nigericin (Figure 1D) and NH_4_Cl (Supplementary Figure S4), the latter having been often used to dissipate NPQ in diatoms (Lavaud et al., 2002a; Ruban et al., 2004; Goss et al., 2006). Both compounds suppressed NPQ in a dose-dependent manner (Figure 1D and Supplementary Figure S4), although high concentrations (i.e. 100 µM nigericin and over 50 mM NH4Cl) were required to substantially inhibit fluorescence quenching (see below).

### The K^+^/H^+^ antiporter KEA3 drives the pH/NPQ relationship in *P. tricornutum*

Having established the crucial role of the pH component of the PMF on *P. tricornutum* NPQ kinetics, we decided to investigate how the PMF is regulated in diatoms, where this phenomenon is still largely ignored. The action mechanism of nigericin (Figure 1D), which is a K^+^/H^+^ exchanger drug, reminds of that of the K^+^/H^+^ antiporter KEA3 (Tsujii et al., 2019) previously suggested to modulate acclimation to light transients in *Arabidopsis thaliana* (Kunz et al., 2014; Armbruster et al., 2014; Wang et al., 2017; Szabò and Spetea, 2017). Based on phylogenetic sequence analysis (Supplementary Figure S5), we identified the Phatr3_J39274 gene as the most likely homologue of KEA3 in *P. tricornutum*. This gene contains typical elements of this family of antiporters: the cation/H^+^ exchange and regulator of K^+^ conductance (RCK) domains (Supplementary Figure S6). Moreover, it harbors an EF-hand, possibly involved in Ca^2+^ binding, which is not found in *Viridiplantae* (Figure S5). Consistent with bioinformatic predictions (Gruber et al., 2015), which pinpoint a plastid targeting signal peptide, the gene product fused to eGFP localized in the chloroplast of *P. tricornutum* (Figure 2A). To study the function of the Phatr3_J39274 gene-encoded protein, we generated knockout (KO) mutants using a CRISPR-Cas9 approach (Allorent et al., 2018). We selected two clones (*kea3-1* and *kea3-2*) containing truncated proteins without the catalytic transmembrane domain (Supplementary Figure S6 A and B). The PtKEA3 protein, which migrated as a double band in SDS-PAGE gels under standard conditions (see below for an explanation) was no longer detectable in the KO strains (Figure 2B). We complemented *kea3-1* with PtKEA3 fused to eGFP under the control of the FCP promoter (Siaut et al., 2007; Falciatore et al., 2020). This genotype, named *kea3-1/KEA3-eGFP* (Figure 2B, see also Supplementary Figure S7 for other complemented lines) turned out to be an overexpressor (OE) of PtKEA3 at both transcriptional and translational levels (Figure 2B, 2C), and will be referred to as an OE hereafter.

**Figure 2.**
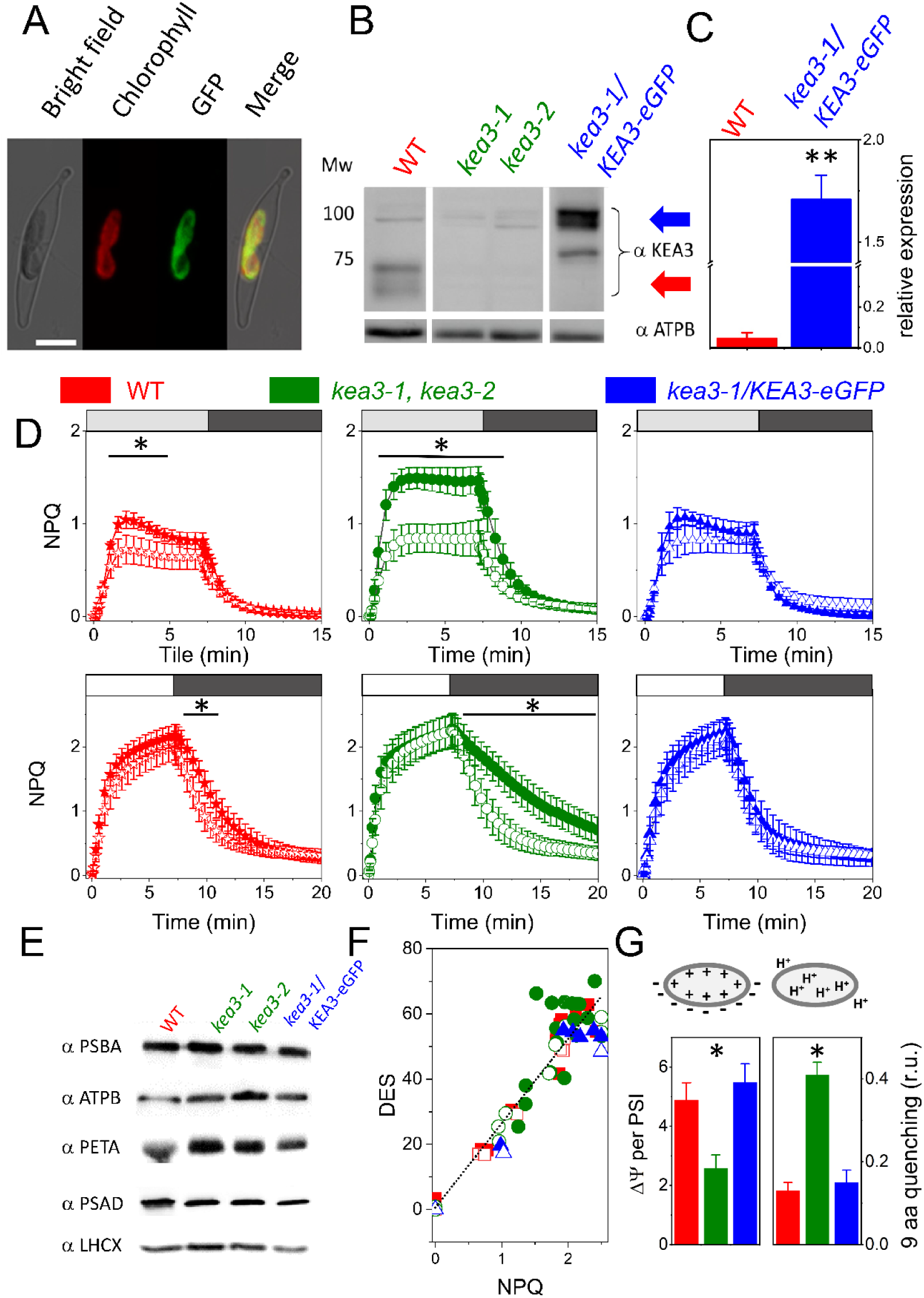
Molecular and physiology phenotypical characterization of KEA3 mutants in *P. tricornutum.* (**A**) Localization of a full-length PtKEA3:eGFP fusion protein expressed in *P. tricornutum*. Bright field: light microscope images; chlorophyll: chlorophyll auto-fluorescence; eGFP: GFP fluorescence; merge: merged channel. Scale bar: 5 μm. (**B**) Immunodetection of KEA3 in total protein extracts from WT (red arrow) and *kea3-1*, *kea3-2* and *kea3-1*/KEA3-eGFP (blue arrow) strains. 30 µg total protein content was loaded per well. Note that the higher MW in the complemented lines is due to the presence of eGFP. Loading control: β subunit of the plastidial ATP synthase complex (ATPB). Representative picture of an experiment repeated 3 times with similar results. (**C**) qPCR quantification of KEA3 mRNA steady state in WT (red) and complemented kea3-1/KEA3-eGFP genotypes (blue). N = 3 mean ± SD. Expression was normalized on three house-keeping genes (RPS, TBP and EF1-a). The asterisks indicate significant differences in expression between the WT and kea3-1/KEA3-eGFP genotypes (P < 0.01). (**D**) NPQ features in WT (red) KO (green) and complemented (blue) genotype under moderate light (ML: 125 µmol photons m^-2^ s^-1^, upper panels) and high light (HL: 450 µmol photons m^-2^ s^-1^, lower panels) in control conditions (solid symbols) and upon addition of 10 µm nigericin (open symbols). White box: HL; light grey box: ML; dark grey box: Low Light (25 µmol photons m^-2^ s^-1^) to facilitate NPQ recovery. ML: N = 6 mean ± SD. HL: N = 15 mean ± SD. Asterisks indicate significant differences in NPQ between control and nigericin treated samples (P < 0.05) (**E**) Western blot analysis of photosynthetic complexes in WT and mutant genotypes. Total protein extracts were analyzed by immunodetection using specific antisera. PSBA: Photosystem II D1 protein 25 µg protein loaded; Cyt f: cyt *b_6_f* complex PETA protein, 50 µg protein loaded; PSAD: Photosystem I subunit, 20 µg protein loaded; LHCX, 20 µg; ATPB: ß subunit of ATPase, 20 µg protein loaded, as loading control. Representative picture of an experiment repeated 3 times with similar results (**F**) Relationship between NPQ capacity and DES in the different genotypes. Same color code as in panels C and D. N = 5 mean ± SD. Solid symbols: control; open symbols: nigericin 10 µM. (**G**) Estimates of the two components of the transthylakoid Proton Motive Force (PMF): the electric potential via the ECS signal (left; N = 6 mean ± SD) and the proton gradient via the quenching of 9-aminoacridine (9-aa) fluorescence (right; N = 4 mean ± SD). Raw data shown in Supplementary Figure S11. Asterisks indicate significant differences in ΔΨ or ΔpH indicative signals between the *kea3-*1, *kea3-2* (pooled data from the two lines), the WT and the *kea3-1/KEA3-eGFP* genotypes (P < 0.05).

The KEA3 KO mutant cells showed an altered NPQ response at moderate light (ML, 125 µmol photons m^-2^ s^-1^), where fluorescence quenching was higher than in the WT and complemented lines (Figure 2D, Supplementary Figure S7, Supplementary Figure S8, Supplementary Figure S9). Addition of nigericin restored WT-like NPQ values in the KO mutants, while being almost without effect in the other strains (Figure 2D). Differences in the NPQ extent and nigericin sensitivity were lost in HL (above 450 µmol photons m^-2^ s^-1^; Figure 2D and Supplementary Figure S9). However, KEA3 KO strains displayed faster kinetics of NPQ onset under HL, while in low light, NPQ relaxation was slower in KO strains than in WT and OE cells (Supplementary Figure S10), in line with earlier results in plants (Armbruster et al., 2014). The NPQ phenotype of KO mutants was not due to changes in the accumulation of specific photosynthetic complexes, for which we could detect comparable levels of representative proteins in all strains (Figure 2E). This also included the main and constitutive NPQ effector in *P. tricornutum* (LHCX1; (Bailleul et al., 2010)), hence excluding a different accumulation of this protein as the cause of the change in NPQ capacity observed in the mutants. The Stern-Volmer relationship between the NPQ extent and de-epoxidation state (DES, Figure 2F) was also similar in all genotypes, both in the absence and presence of nigericin. This result, already reported in the case of WT *P. tricornutum* cells treated with NH_4_Cl (Goss et al., 2006), suggests that PtKEA3 modulates the relationship between NPQ and the PMF without altering the S-V relationship between fluorescence quenching and the XC.

Overall, these data indicate that: *i.* nigericin is effective in modulating *P. tricornutum* NPQ in vivo. *ii.* its effect is enhanced upon inactivation of the PtKEA3 H^+^/K^+^ antiporter. *iii.* overexpressing PtKEA3 makes NPQ nigericin insensitive. Based on previous observations in plants (Kunz et al., 2014; Armbruster et al., 2014; Wang et al., 2017; Galvis et al., 2020), a plausible explanation for the different effect of nigericin on NPQ in WT and mutants would be that the ΔpH is enhanced upon removal of the KEA3 antiporter. The consequently increased ΔpH would trigger a larger, nigericin-sensitive NPQ in the KO mutants. We tested this possibility by comparing the two components of the PMF (i.e. the electric field – ΔΨ – and the proton gradient – ΔpH (Witt, 1979)) *in vivo*. We measured changes in the ΔΨ with the ECS signal, a modification of the absorption spectrum of specific pigments that senses the transmembrane electric field ((Witt, 1979), Figure 2G and Supplementary Figure S11; see also Methods). In parallel, we implemented the use of the fluorescent probe 9-aminoacridine (9-aa) to estimate changes in the ΔpH (Schuldiner et al., 1972) in living cells (Figure 2G and Supplementary Figure S11; see Methods). We observed complementary changes in the two PMF components in the different genotypes. In particular, the increased ΔpH in light-exposed KEA3 KO cells was compensated by a similar decrease in the ΔΨ component (Figure 2G), while the KEA3 OE mutant displayed a WT-like partitioning of the PMF.

### KEA3 controls NPQ by modulating XC dynamics in diatoms

Although the ΔΨ/ΔpH changes may account for the mutants NPQ phenotype in ML, they do not explain why NPQ becomes similar in all genotypes in HL. We hypothesize that although the proton fluxes might be different in the different genotypes, the luminal pH may become acidic enough to trigger similar NPQ responses in HL. We verified this possibility adapting a computational model recapitulating NPQ features in plants (Matuszyńska et al., 2016) to diatoms (Figure 3A, see Supplemental Information). To simulate the activity of PtKEA3, we introduced a partitioning of the PMF between its ΔΨ and ΔpH components. While both can contribute to ATP synthesis (Witt, 1979), only the latter is able to modulate NPQ. We simulated the consequences of the KEA3 KO and OE mutations by modifying the ratio (K_KEA3_) between the ΔΨ and ΔpH according to the changes in the two PMF components observed in the different genotypes (Figure 2G). This led us to satisfactorily recapitulate the main experimental features of NPQ in diatoms, including the higher NPQ extent in KO mutants under ML (Figure 3B) and the light dependency of NPQ (Figure 3C) in the three genotypes, where NPQ became comparable in HL. Based on these results, we conclude that: *i.* steady state NPQ is regulated by pH under different light conditions. *ii.* this regulation mainly depends on changes in the ΔΨ to ΔpH ratio. Based on the mutant phenotypes found in this study, *iii.* KEA3 appears to be a key regulator of the ΔpH/XC/NPQ relationship in *P. tricornutum*.

**Figure 3.**
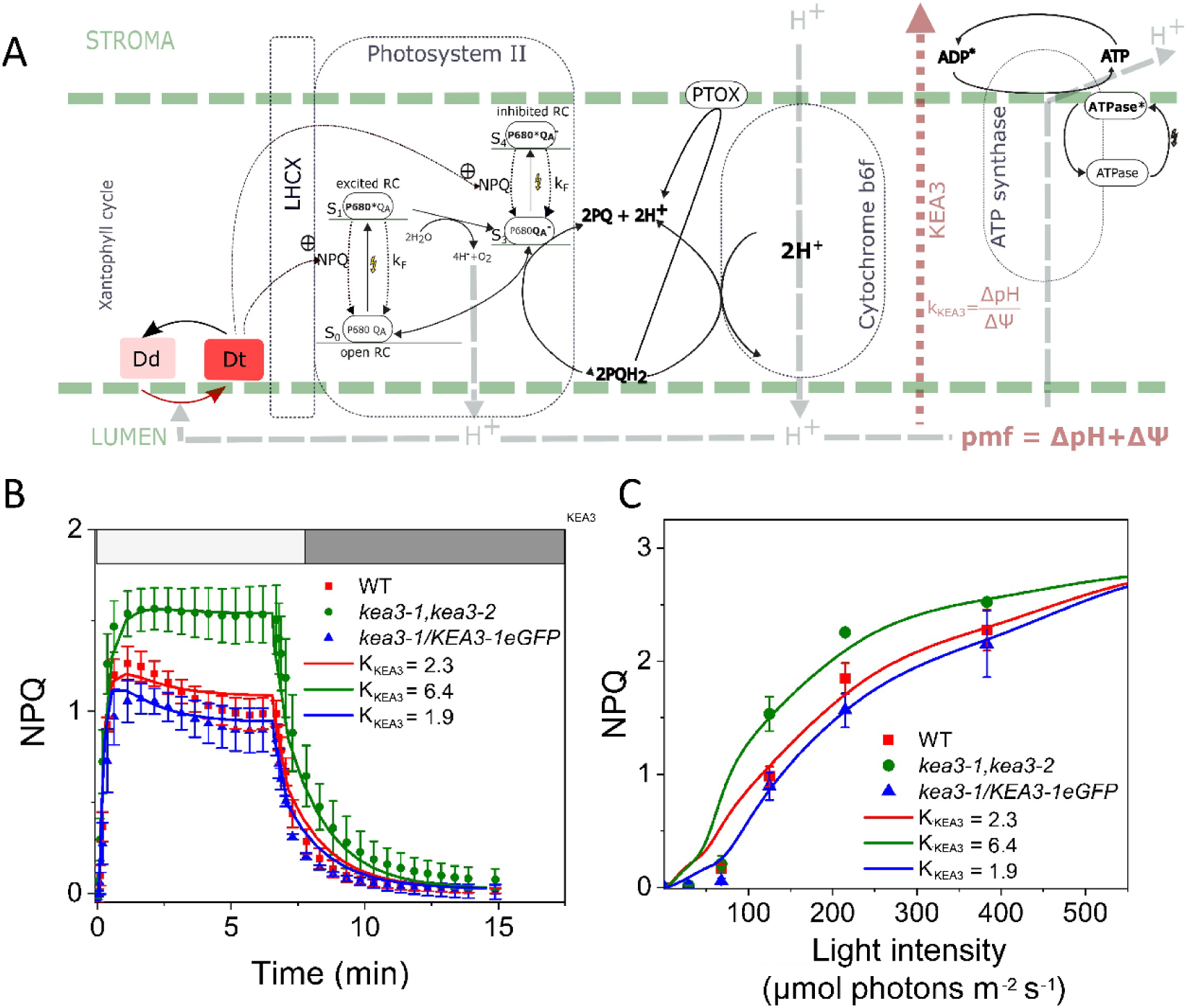
Mathematical model recapitulating NPQ features of diatoms. **(A)** General features of the computational model used to describe diatoms NPQ. The model is adapted from a plant NPQ model (Matuszyńska et al., 2016) with a few modifications (see also methods) including: *i.* a more neutral pK for diadinoxanthin de-epoxidase (6.3) than for violaxanthin de-epoxidase). *ii*. a partitioning of the PMF between the ΔΨ and the ΔH is included via the K_KEA3_ parameter (K_KEA3_ = ΔpH/ΔΨ). **(B)** Simulations of NPQ kinetics on WT and mutants at ML (Raw data from Figure 2D (symbols) were simulated (lines) considering changes in the ΔpH/ΔΨ ratio calculated based on the data of Figure 2G. **(C)** Light dependency of NPQ in the different genotypes. Raw data from Supplementary Figure S9 (symbols) were simulated (lines) considering the same changes in the ΔpH/ΔΨ as in panel B. **(B)**

### Regulation of KEA3 and diatom physiological responses

Having assessed the role of KEA3 on *P. tricornutum* NPQ, we looked for a possible growth phenotype of KEA3 KO mutant cells under different light conditions. Mutants did not display specific growth impairments under different light conditions including ML (Supplementary Table S1), in which we observed differences in the NPQ phenotype at steady-state (Figure 2D). Growing cells in f/2 medium with a 14hL/10hD photoperiod, as already used in other studies (e.g. (Giovagnetti and Ruban, 2017)), substantially modified the NPQ responses of *P. tricornutum* (Supplementary Table S2), without changing the mutant growth capacity. Cells grown in these conditions became more sensitive to inhibition of NPQ by protonophores (especially NH_4_Cl; Supplementary Table 2). When grown under intermittent light (IL), all genotypes displayed a reduced growth rate compared to the 14hL/10hD or the 12hL/12hD photoperiods (Supplementary Table S1), in line with the substantial (seven fold) decrease in the daily dose of received photons (Lavaud et al., 2002c; Giovagnetti and Ruban, 2017). At this light, however, *KEA3* mutants showed a growth retard when compared to the WT and OE lines (Supplementary Table S1), possibly due to very high levels of NPQ formed (> 8), which could compete with PSII light-harvesting capacity for available photons (Giovagnetti and Ruban, 2017; Buck et al., 2019).

Besides the role of KEA3 in modulating transient NPQ responses and acclimation to different light regimes, which is in line with previous results in plants (e.g. (Armbruster et al., 2016; Li et al., 2021)), we found a unique feature of KEA3 in *P. tricornutum*, i.e. a possible role in diel modulation of NPQ. This is suggested by the observation that the NPQ amplitude and sensitivity to nigericin increases at the end of the light phase in WT cells (Figure 4A), while remaining nigericin sensitive and insensitive, respectively, in KO and OE mutants throughout the light phase of the day. Analysis of available transcriptomic data suggests a rationale for this observation, showing that KEA3 transcripts decreased significantly at the end of the light phase in WT cells (Chauton et al., 2013). A decreased KEA3 accumulation in the WT at the end of the light phase should make these cells more alike KO mutants, explaining their enhanced nigericin sensitivity. We confirmed a decrease in PtKEA3 transcripts in WT cells from the beginning to the end of the light phase (Figure 4B). However, we could not detect any change in the PtKEA3 accumulation between samples collected either in the morning or in the evening (Figure 4C), suggesting that changes in the nigericin sensitivity of NPQ cannot be simply explained by changes in PtKEA3 abundance. Recent work has suggested that the plant KEA3 is regulated through its dimerization through the C terminus RCK domain (Uflewski et al., 2021), in line with earlier findings in the bacterial K^+^ efflux system KefC (Roosild et al., 2009). We tested this possibility and found the same oligomerization of KEA3 in thylakoids isolated from cells harvested either in the morning or afternoon (Figure 4D).

**Figure 4.**
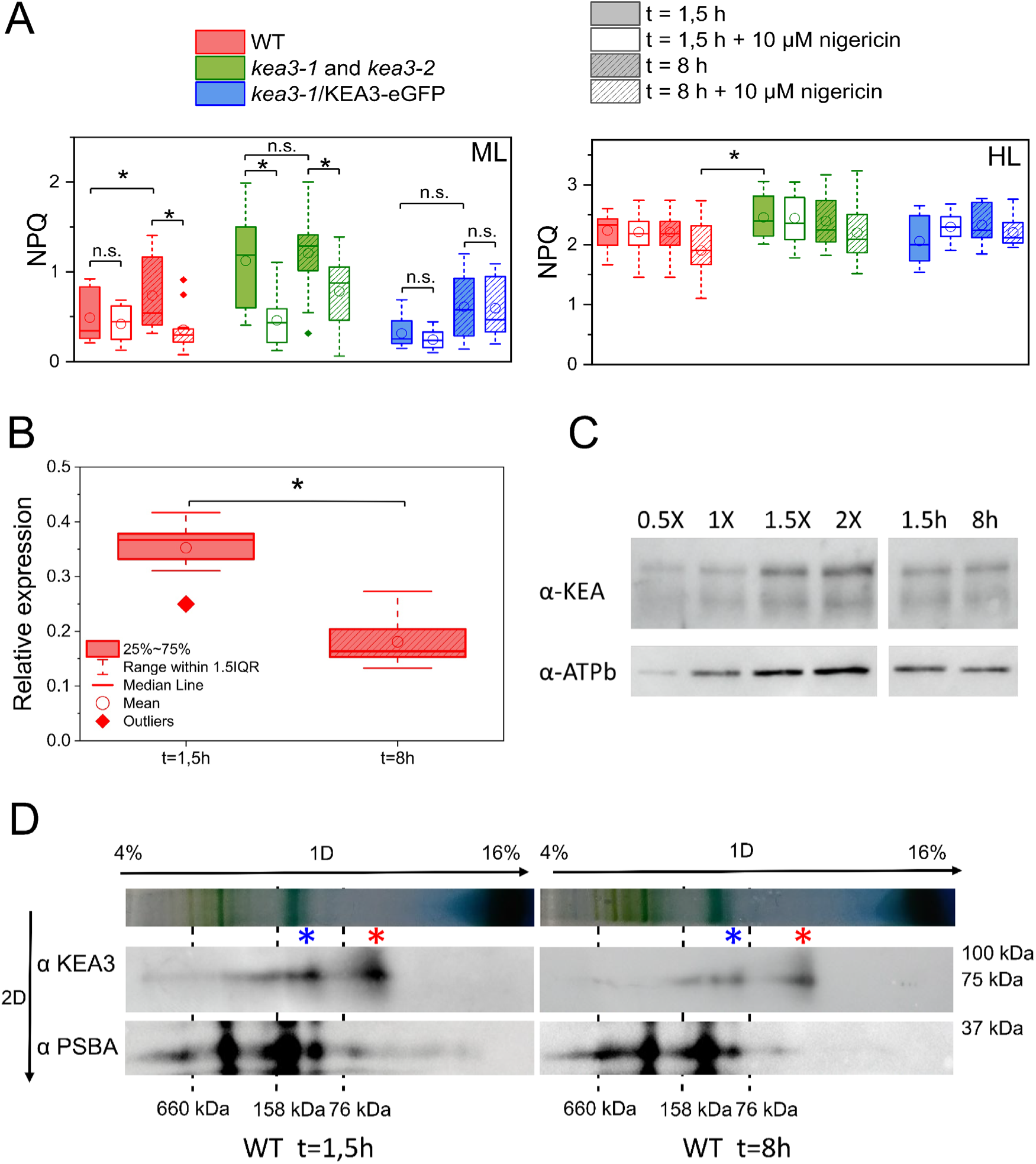
Role of KEA3 in promoting diel variations in the ΔpH sensitivity of NPQ in *P. tricornutum.* Cells were grown in a 12:12 photoperiod and harvested either 1.5 h or 8 h after the onset of illumination. (**A**) Variation in NPQ amplitude and nigericin sensitivity measured in WT, *kea3-1* and *kea3-2* (pooled data from the two lines) and *kea3-1/*KEA3-eGFP cells after 1.5 h (solid, open) and 8 h (dashed) of exposure to ML (125 µmol photons m^-2^ s^-1^, left) and HL (1200 µmol photons m^-2^ s^-1^, right). N = 7-16. (**B**) Quantification of KEA3 mRNA accumulation by qPCR after 1.5 h (solid) and 8 h (dashed) of light exposure to ML light (inset) in WT cells. N = 3. The asterisk indicates significant differences in KEA3 expression levels (P < 0.05). (**C**) Immunodetection of KEA3 protein in WT cells steady state in the thylakoids after 1.5 h and 8 h of illumination. The sensitivity of the Western Blot blot using progressive dilutions of the sample is shown. Representative picture of an experiment repeated 3 times with similar results. ATP-B immunodetection was used as loading control. (**D**) Oligomerization state of KEA 3 in WT cells harvested 1.5 h or 8 h after the onset of illumination during the day. Thylakoid membranes of WT were solubilized with α-DM and loaded on 4-16% Acrylamide BN gel (top arrows). A mix of Thyroglobuline (660 kDa), Aldolase (158 kDa) and Conalbumine (76 kDa) was used as molecular weight markers. PtKEA3 was detected on 2D denaturizing condition using α-KEA3 antiserum and α-PSBA antiserum was used as a running control. Molecular weight are reported on the right. Red asterisk: PtKEA3 monomer; blue asterisk: PtKEA3 dimer. Representative picture of an experiment repeated 3 times with similar results.

Overall these data raise the question of how the function of this protein is regulated in diatoms. The *KEA3* gene has a peculiar EF hand domain in *P. tricornutum*, similarly to other genes involved in cell energetic metabolism (Prihoda et al., 2012). This Ca^2+^ binding domain is located close to the RCK domain (Supplementary Figure S6), i.e. in a position where it could modulate the activity of the antiporter (Wang et al., 2017; Galvis et al., 2020). We therefore investigated a possible role of Ca^2+^ on *P. tricornutum* KEA3. As mentioned above, two KEA3 bands are observed by immunodetection after separation by SDS-PAGE, both in the WT and in OE lines (Figure 2B, Figure 5A). We further investigated the nature of these bands, and their possible relationship to Ca^2+^ binding, using two OE clones, exploiting the fact that signals were larger and easier to analyze in these genotypes. We found that addition of 1mM CaCl_2_ to the gel led to the appearance of only a single band, corresponding to the lower band of the control (blue asterisk) (Figure 5B). Conversely, the Ca^2+^-chelator EGTA (10 mM) had an opposite effect, increasing the intensity of the upper (orange asterisk) band (Figure 5C). Replacement of CaCl_2_ with 1 mM MgCl_2_ had no consequences on the protein migration compared to standard conditions (Figure 5D). These findings indicate that *i.* Ca^2+^ binds to PtKEA3 and alters the protein mobility in the gel in agreement with previous findings with other Ca^2+^ binding proteins bearing EF-hand motifs (Ishitani et al., 2000; Kamthan et al., 2015) *ii.* the effect of this ion is specific (as indicated by the lack of effect of MgCl_2_) and can be reversed to some extent by Ca^2+^ chelation. Thus, the *in vitro* binding of PtKEA3 to Ca^2+^ suggests a possible role of this ion in controlling protein conformational changes and therefore activity, as previously reported in the case of other EF-hand-bearing Ca^2+^ binding proteins (e.g. (Ishitani et al., 2000; Kamthan et al., 2015) ). We further explored this hypothesis generating *P. tricornutum* KEA3 mutants lacking the EF-hand motif (Figure S6A). Although these cells still express the KEA3 gene to WT-like levels (Figure 5E), they displayed a much higher and nigericin sensitive NPQ than the WT in ML (Figure 5F), i.e. similar characteristics as the KEA3 KO mutants (Figure 4A). This finding suggests that the antiporter was inactive in these genotypes. We conclude that Ca^2+^ modulates KEA3 activity in *P. tricornutum* via its EF-hand in diatoms, unlike plant KEA3 (Uflewski et al., 2021). Based on previously reported daily changes in the cellular Ca^2+^ concentration during the day in plants (e.g. (Love et al., 2004)) it is tempting to suggest that intracellular variations/fluctuations in the concentration of this ion may be involved in the diurnal changes in KEA3 activity/NPQ observed in *P. tricornutum*.

**Figure 5.**
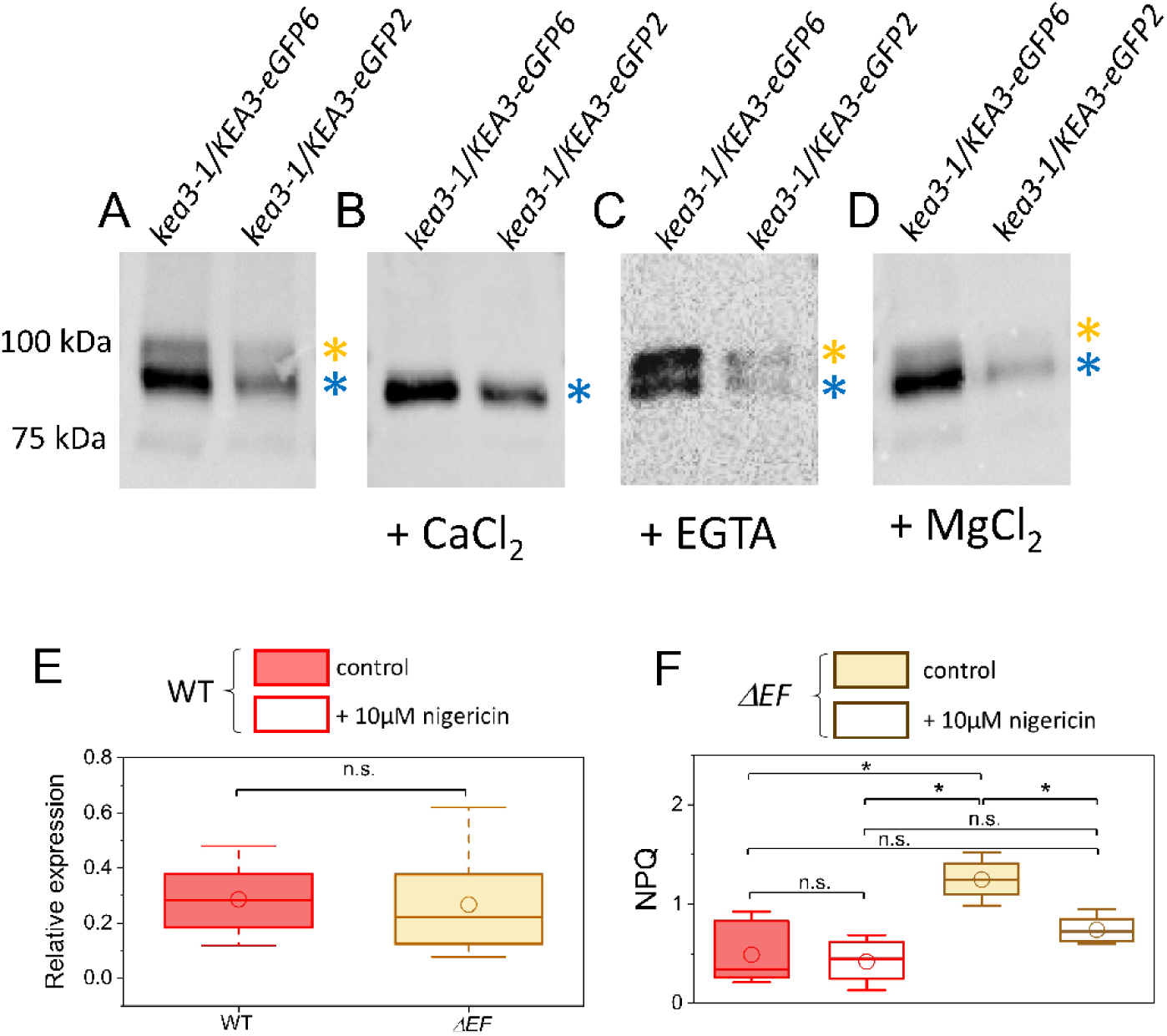
Ca^2+^ binding controls KEA3 activity in *P. tricornutum* through an EF-hand motif. Immunodetection of KEA3 of total protein extracts from two different kea3-1/KEA3-eGFP overexpressing strains (see Supplementary Figure S7). The polypeptides were separated by SDS-PAGE and revealed with an antibody directed against eGFP (Miltenyi biotech, US). (**A**) control conditions: kea3-1/KEA3-eGFP appears as a doublet (orange and blue asterisks). (**B**) In the presence of 1mM CaCl_2_, only a single band is visible. (**C**) The Ca^2+^-chelator EGTA (10 mM) has an opposite effect, increasing the intensity of the upper (orange asterisk) band. (**D**) replacement of CaCl_2_ with 1 mM MgCl_2_ has no consequences on the polypeptide migration. (**E**) Quantification of KEA3 mRNA accumulation by qPCR in the WT (red) and in KEA mutants lacking the EF calcium binding motif (*ΔEF-1* and *ΔEF-2*, pooled data from the two lines). N = 3. (**F**) NPQ amplitude and nigericin sensitivity measured in the WT and in *ΔEF* lines (*ΔEF-1* and *ΔEF-2*, pooled data from the two lines). N = 12-27. Asterisks indicate significant differences in NPQ (P < 0.05). *ΔEF* lines are described in Supplementary Figure S6. Statistical parameters concerning panels (A), (B) (E) and (F) are indicated in panel B.

## Discussion

NPQ has been extensively studied in diatoms, not only because of the ecological relevance of this microorganism, but also because of its peculiar features when compared to plants. Diatoms harbor specific NPQ protein effectors (LHCX) as well as a peculiar XC (reviewed in (Falciatore et al., 2020)). The amplitude of NPQ in *P. tricornutum* largely exceeds the one observed in plants and green microalgae (Lavaud et al., 2002b; Giovagnetti and Ruban, 2017; Taddei et al., 2018). Diatoms have been proposed to develop different quenching sites depending on environmental conditions (Chukhutsina et al., 2014; Giovagnetti and Ruban, 2017; Taddei et al., 2018). Despite significant progress in understanding the mechanism of diatom NPQ process, the relationship between fluorescence quenching and thylakoid membrane energizations, which is a basic feature of plant NPQ (Wraight and Crofts, 1970), is still debated and not fully understood (Goss et al., 2006; Lavaud and Kroth, 2006).

In *Viridiplantae*, the proton gradient activates both the XC and NPQ effectors (PSBS in vascular plants, LHCSR and PSBS in mosses and green algae (Niyogi and Truong, 2013; Giovagnetti and Ruban, 2018)). Due to their different apparent pKa, these PMF targets lead to complex NPQ responses to the proton gradient (Zaks et al., 2012; Matuszyńska et al., 2016). In diatoms, the pH dependence of NPQ reported in Figure 1 reminds that of the diatom diadinoxanthin de-epoxidase (Jakob et al., 2001) consistent with the notion that this enzyme is the main target of the PMF. However, we cannot exclude a pH regulation of LHCXs because the pH dependency of NPQ found in *P. tricornutum* (Figure 1) is also similar to that reported in *C. reinhardtii* (Tian et al., 2019), where LHCSR3 is instead the main target of the ΔpH regulation (Ballottari et al., 2016). These findings suggest that the pH dependency of the LHCX protonation and XC is either the same in diatoms or it is dominated by the de-epoxidase activity, likely due to a different pH regulation (if any) of the function of LHCXs relative to LHCSRs (Shinkle et al., 2010). In line with the second possibility, we observed that the pH dependency of NPQ is the same in Pt1 (our reference ecotype) and the Pt4 ecotype, although the latter displays significantly reduced, constitutive levels of LHCX1 and thus a lower extent of NPQ (Fig. S12; (Bailleul et al., 2010)). Conversely, one would expect different pH profiles in the two ecotypes if LHCX1 and the diadinoxanthin de-epoxidase respond to pH with different profiles, because of the diminished contribution of LHCX1 in Pt4.

Multiple proteins (e.g. TPK3, KEA3, CLCe, VCCN1, PHT4;1, PAM71/CCHA1) have been invoked as modulators of the partitioning between the ΔΨ and ΔpH components of the PMF in plants, acting as ion channels or antiporters (reviews by (Szabò and Spetea, 2017; Li et al., 2021)). In diatoms, knocking out the KEA3 gene strongly affects NPQ transients and steady state levels in both intermittent and continuous light (at different intensities; Supplementary Figure S8 and Supplementary Table S1). KEA3 is therefore a key regulator of NPQ in *P. tricornutum* during transients, in line with earlier conclusions in plants (Armbruster et al., 2014; Kunz et al., 2014). Our model simulations (Figure 3) extend this role in the model diatom studied here, since we can reproduce most of the main features of *P. tricornutum* NPQ in steady state conditions assuming the existence of a simple regulatory circuit of NPQ with one elicitor (the ΔpH component of the PMF), a master PMF regulator (KEA3) and one target (most likely the XC, Figure 3A). On the other hand, NPQ transients (and in particular the slower NPQ relaxation in KO strains) are not well reproduced by these simulations (Figure 3B), suggesting that other effectors may also fine tune NPQ responses in *P. tricornutum* during transients, as recently suggested in plants (Li et al., 2021). Consistent with this possibility, several of the putative PMF regulators identified in plants (e.g. TPK3, CLCe, VCCN1) are also found in diatoms genomes (Marchand et al., 2018). Diatoms display a panoply of NPQ responses in terms of extent, regulation and, likely, quenching sites. This flexibility likely stems from the presence of numerous LHCX isoforms (e.g. 4 in *P. tricornutum* and up to 11 in *Fragilariopsis cylindrus*; (Taddei et al., 2016; Mock et al., 2017), which catalyze responses to low/moderate light (LHCX1), high light (LHCX2, LHCX3), blue light (LLHCX1-3, (Coesel et al., 2009)), iron (LHCX2), nitrogen limitation (LHCX3, LHCX4) as well as the onset of different quenching sites depending on light conditions (Chukhutsina et al., 2014; Taddei et al., 2018).

The existence of mechanisms linking the PMF to chloroplast changes may provide an additional layer of regulation. The PtKEA3 C-terminus contains a conserved regulatory domain which should regulate its activity upon binding of nucleotides (ATP, ADP, AMP, and NAD(P)H (Roosild et al., 2002; Cao et al., 2013).In plants, the topology of this antiporter is still discussed (Armbruster et al., 2016; Wang et al., 2017) but a stromal location of the terminal domain is possible, and would allow regulation of the antiporter by stromal nucleotides. It has been hypothesized that light mediated changes in the ATP/ADP and/or NADPH/NADP^+^ ratios might participate in the modulation of the PMF, via their effect on KEA3 (Galvis et al., 2020) and therefore of light acclimation itself via NPQ (Uflewski et al., 2021). The fact that in diatoms, NADPH and ascorbate are mandatory co-factors for the diatoxanthin epoxidase and diadinoxanthin de-epoxidase, respectively (Goss et al., 2006; Blommaert et al., 2021), may provide an additional link between the redox poise of the stroma and NPQ, adding an additional dimension to the role of PtKEA3 in NPQ control.

Our data are in principle consistent with such possibilities. However, we reveal here the existence of a peculiar regulation (the Ca^2+^ sensitivity of PtKEA3 provided by its EF hand domain shared with other diatoms’ KEA3 homologs), which is seen in the model diatom *P. tricornutum,* and likely provides a link between the PMF and environmental changes, i.e. the diel regulation of NPQ reported here. This regulation could reflect the relevant role of Ca^2+^ in mediating responses to fluid motion, osmotic stress, and iron, i.e. environmental challenges experienced by phytoplankton in turbulent waters (Falciatore et al., 2000), while likely being being less important for sessile plants specific Previous work has already shown a linkage between intracellular Ca^2+^ and NPQ in green microalgae (Petroutsos et al., 2011). However, the mechanism in green algae differs from the one reported here: while in *P. tricornutum* Ca^2+^ seems to affect the PMF, i.e. the ‘source’ of NPQ, via KEA3, this ion modulates the accumulation of the LHCSR3 protein, i.e. the NPQ ‘effector’ in *C. reinhardtii*.

These mechanisms, which deserve to be considered in a more refined model of NPQ regulation, may further contribute to the required functional diversity/flexibility of light-harvesting regulation for diatoms to thrive in different aquatic environments, where light acclimation is often a major determinant for growth and survival (Lin et al., 2016).

## Materials and Methods

### Cells growth

*P. tricornutum* Pt1 and Pt4 cells were grown in ESAW (Enriched Seawater Artificial Water, (Berges et al., 2001)) supplemented with NaNO_3_ 468 mg L^-1^ and NaH_2_PO_4_.H_2_O 30 mg L^-1^, at 20°C, a light intensity of 40 µmol photons m^-2^ s^-1^, a 12h light-12h dark photoperiod and shaking at 100 rpm. Alternatively, *P. tricornutum* Pt1 cultures were grown in sterile artificial seawater f/2 medium (GUILLARD and RYTHER, 1962) at 18 °C, and shaking at 100 rpm with cells exposed either to a 14 h light/10 h dark photoperiod (40 μmol photons m^−2^ s^−1^) or intermittent light (IL, 40 μmol photons m^−2^ s^−1^, with cycles of 5 min light/55 min dark) (Giovagnetti and Ruban, 2017).

### Identification of the KEA3 homologue in *Phaeodactylum tricornutum* and bioinformatics analysis

The gene Phatr3_J39274 was identified as the most likely homologue of AtKEA3 (AtKEA3, At4g04850) by BLAST on the genome of *P. tricornutum*. Alignment and phylogenetic reconstructions were performed using the function “build” of ETE3 v3.1.1 (Huerta-Cepas et al., 2016) as implemented on GenomeNet (https://www.genome.jp/tools/ete/). Alignment was performed with MAFFT v6.861b with the default options (Katoh and Standley, 2013). ML tree was inferred using PhyML v20160115 ran with model JTT and parameters: --alpha e -o tlr -- nclasses 4 --pinv e --bootstrap -2 -f m (Guindon et al., 2010). The protein sequences used in this work are available in Supplementary Table S5.

The domains of PtKEA3 were identified using the following web tools: InterPro (https://www.ebi.ac.uk/interpro/), ASAFind (https://rocaplab.ocean.washington.edu/tools/asafind/) and SignalP 4.1 (http://www.cbs.dtu.dk/services/SignalP-4.1/) (Gruber et al., 2015; Bendtsen et al., 2004).

### Construction of KO-mutants and mutant complementation by biolistic transformation

All primers used in this work are available in Supplementary Table S4. Target sites for CRISPR-Cas9 were identified within the Phatr3_J39274 gene using the PhytoCRISP-Ex web tool (https://www.phytocrispex.biologie.ens.fr/CRISP-Ex/). Adapters were designed according to the protocol of (Allorent et al., 2018). The gene coding for zeocin resistance was inserted into the plasmid pKSdiaCaS10_sgRNA (Addgene, plasmid #74923) by Gibson cloning (Gibson et al., 2009) to create the plasmid pKSdiaCAs9_sgRNA_ZeoR. The adapters were then inserted into the plasmid pKSdiaCas9_sgRNA_ZeoR. The resulting plasmids were coated on tungsten beads and used for biolistic transformation of the cells with the PDS-100/He system (Bio-Rad, 1672257). DNA from zeocin (100 µg/mL final) resistant transformants was extracted, amplified by PCR and sequenced for mutant identification.

To complement KO mutants, Genomic DNA was extracted from wild-type Pt1 cells and the gene Phatr3_J39274 was amplified by PCR using the Phusion^TM^ Polymerase (Thermo Fisher) (see table below). The CDS was fused to an eGFP encoding DNA sequence at the C-terminal and inserted into a vector containing a gene for blasticidin resistance under control of the Fcp promoter. The plasmid was transformed into *E. coli* Top 10 cells. The cells were lysed and the harvested plasmids were used for biolistic transformation of the mutant *kea3-1* using the same procedure as for the construction of KO-mutants. Transformants were selected on ESAW supplemented with zeocin (100 µg/mL) and blasticidin (12 µg/mL).

### qPCR

Total RNA was extracted from cell cultures of WT KEA3 KO and OE genotypes as previously described (Siaut et al., 2007). Briefly, cells were collected after having been rinsed twice with adapted PBS buffer (15mM Na_2_HPO_4_, 4.5mM NaH_2_PO_4_, 490mM NaCl, 0.1mM EGTA). Then cells were re-suspended in TRI Reagent (Sigma) and then centrifuged under 10000 rpm to remove the unbroken cells and genomic DNA. RNA in suspensions were extracted by chloroform and following by isopropanol.

cDNA was synthesized from 1 µg of purified RNA of each samples using cDNA high yield synthesis kit (SuperScript IV VILO, Thermo Fisher Scientific). Quantitative PCR was performed with following primers and SYBR green kit (PowerSYBP Green PCR Master Mix, Thermo Fisher Scientific) in a total volume of 10 µL. qPCR amplification reactions were run and analyzed on CFX-Connect with following cycling conditions: 95°C 10min, 40 cycles of 95°C 10s and 60°C 40s, 95°C 10s, melt Curve 65°C to 95°C, increment 0.5°C followed by plate read.

### Localization by confocal imaging

For the localization analysis, the cDNA sequence of Phatr3_J39274 was amplified using the listed primers with *P. tricornutum* cDNA as a template, and then inserted into the pPhaT1-linker-eGFP vector according to the manufacturer’s instructions (C115, Vazyme Biotech, China). To obtain the vector pPhaT1-linker-eGFP, a linker (GGACCTAGGGGAGGAGGAGGAGGA) was inserted before the eGFP coding sequence and then linker-eGFP was cloned into the Xba I-Hind III sites of pPha-T1. Inserts of all constructs were sequenced to confirm that no unintended nucleotide changes were present. Vectors were introduced into *P. tricornutum* by electroporation as previously described (Zhang and Hu, 2014). The transgenic algae were visualized under a Leica TCS SP8 laser scanning confocal microscope. Chlorophyll autofluorescence and GFP fluorescence were excited at 488 nm and were detected with 630–690 nm and 500–550 nm, respectively.

### Recombinant protein expression and purification. Production of α--KEA3 antibody

The KEA3 C-terminal cDNA was amplified by PCR from *P. tricornutum* genomic DNA, using the primers listed below. The PCR product was cloned into a pet28b(+) vector by Gibson assembly (Gibson et al., 2009). The construct was named KEA3 Cterm pet28 b(+). The protein was fused with a N-terminal His-tag.

*E. coli* Rosetta 2 competent cells were transformed with KEA3 Cterm pet28 b(+) construct and grown at 37°C in the presence of Chloramphenicol (34 μg.mL-1) and Kanamycin (50 μg mL-1) until absorption at 600 nm reached an optical density value of 0.6. Upon addition of IPTG (0.4 mM) growth was continued at 18 °C for 15 h. Bacteria were pelleted, suspended in buffer A (25mM Tris pH 8.0, 300mM NaCl, 1mM TCEP, 5% glycerol, 1 mM benzamidine, 5 mM ε-aminocaproic acid) and lysed by sonication. Insoluble material was removed by centrifugation (15,000 g, 20 min, 4°C), and soluble proteins were recovered. The protein extract was loaded onto a Ni-NTA column. After washing with buffer A supplemented with 50 mM imidazole, the recombinant protein was eluted in buffer A supplemented with 300 mM imidazole. The pure fractions of KEA3 Cterminal were pooled, frozen in liquid N2 and stored at -80 °C.

For antibody production, 2 mg of pure KEA3 C-terminal fraction were loaded on a large SDS-PAGE gel containing 12% acrylamide. After a slow migration overnight, the gel was stained by a solution of 4M sodium acetate, the band containing the recombinant protein was cut. Proteins present in this band were electroeluted in 50 mM NH_4_CO_3_ 0.1 % SDS. This sample was frozen in liquid nitrogen and freeze-dried. The powder was resuspended in 1 mL of PBS.

Antibodies from guinea pig immunization with this purified protein product were obtained from Charles River biologics (USA).

### Protein isolation and immunoblot analyses

Total proteins were isolated from cell cultures in mid-exponential phase in Hepes 50mM supplemented with EDTA-free protein inhibitor cocktail (Roche). Cell walls were broken by three 30-s cycles at 10000 rpm in the presence of micro glass beads (200-400 µm diameter Sigma-Aldrich, Precellys, Bertin Technologies, France). The proteins extracts were clarified by centrifugation and proteins were precipitated for 20 minutes at -20°C in 80% (v/v) acetone. After centrifugation, the pellet was solubilized for 5 minutes at 4°C in 50 mM Tris, pH 7.6, 2% SDS supplemented with EDTA-free protein inhibitor cocktail. For KEA3 protein detection, extracts enriched in membrane proteins were prepared as follows. After cells rupture, total extracts were centrifuged (12000 g, 90s, 4°C) and the pellets (containing membranes) were washed twice with Hepes 50 mM, NaCl 50 mM. Finally, membrane proteins were solubilized with Tris-SDS solution as previously described. Samples were prepared according standard protocols for SDS-PAGE. For BN-PAGE: Chloroplasts were isolated as in (Flori et al., 2017). Cells were collected and resuspended in a small volume of isolation buffer (500 mM sorbitol, 50 mM HEPES-KOH pH 8, 6 mM EDTA, 5 mM MgCl_2_, 10 mM KCl, 1 mM MnCl_2_, 1% PVP 40, 0.5% BSA, 0.1% cysteine). Cells were mechanically disrupted with a French press (90 MPa), unbroken cells were discarded by centrifugation (200 g, 5 min) and cellular fractions were collected and resuspended in 2 mL of washing buffer (500 mM sorbitol, 30 mM HEPES-KOH pH 8, 6 mM EDTA, 5 mM MgCl_2_, 10 mM KCl, 1 mM MnCl_2_, 1% PVP 40, 0.1% BSA) and loaded on discontinuous Percoll gradients (10, 20, 30%). After centrifugation (SW41Ti rotor, 10000 g, 35 min) the fraction at 20-30% interface was collected and diluted in the same buffer (without BSA) and further centrifuged (60000 g, 35 min). The final chloroplast pellet was resuspended in a hypotonic buffer (10 mM MOPS, 4 mM MgCl_2_, protease inhibitor cocktail). Thylakoids were collected by centrifugation (10000 g, 10 min) and stored in 25 mM BisTris pH 7, 20% Glycerol at -80°C.

Thylakoids (100 µg of proteins) were solubilized with 0.75% n-Dodecyl-α-D-maltoside (Anatrace) at RT for 20 min. Unsolubilized membranes were then pelleted by centrifugation (20000 g, 20 min). Prior to gel running 1:10 v/v BN-loading buffer (500 mM Aminocaproic acid, 30% Sucrose, 5% Blue Comassie G-250, 100 mM BisTris pH 7) was added to the supernatant. Gels (NativePAGE^TM^, 4-16%, Bis-Tris, Invitrogen) were run as described in (Järvi et al., 2011) (Cathode buffer: 50 mM Tricine, 15 mM BisTris pH 7, 0.01% Blue Comassie G-250; Anode buffer: 50mM BisTris pH 7). A mix of Conalbumine (76 KDa), Aldolase (158 KDa) and Thyroglubuline (660 KDa) was used as molecular weight marker. After running, single lines were excised from the gel and were soaked with 138 mM Tris pH 6.8, 4.3% SDS for 40 min, followed by 1 minute with the same solution supplemented with 20% Glycerol prior loading onto 8.5% Acrylamide SDS-PAGE. For denaturing gels, polypeptides were separated by SDS-PAGE and transferred to a PVDF membrane for immunodetection following standard protocols. All antibodies were diluted in TBS-Tween 0.1% (v/v) at the following dilutions: α-ATPB (Agrisera, AS05 085) 1:10000, α-PsaB (Agrisera, AS10 695), α-PSBA (Agrisera, AS10 704), α-PETA (Agrisera, AS06 119), α-LhcX (gift of G. Peers, Colorado State University, USA; 1:25000) α –GFP (Miltenyi biotech130-091-833) and custom-made α-KEA3 (1:2000).

To test the consequences of Ca^2+^ on KEA3 migration in SDS-PAGE gels, acrylamide gels were supplemented with 10 mM EGTA, 1 mM CaCl_2_ or 1 mM MgCl_2_. In these cases, the Western blot running buffers were: 0.025 M Tris, 0.192 M glycine, 0.1 % SDS, 10 mM EGTA in the anode and cathode buffer for EGTA blots; 0.025 M Tris, 0.192 M glycine, 0.1 % SDS, 1 mM MgCl_2_ for MgCl_2_ blots; 0.025 M Tris, 0.192 M glycine, 0.1% SDS in the cathode buffer and 0.025 M Tris, 0.192 M glycine, 1 mM CaCl_2_ in the anode buffer for CaCl_2_ blots.

### Chlorophyll fluorescence measurements and NPQ induction in the dark by pH shift

To image chlorophyll fluorescence, 200 µL of cell suspension were placed on wells of a 96-well plate at a cell concentration of 2.10^6^ cells mL^-1^. If needed, nigericin (Nig) or ammonium chloride were added to the cell suspension at this stage. Cells were dark-adapted for 10 min prior to measurements. Wells were imaged for fluorescence emission using a Speedzen III fluorescence imaging setup (JBeam Bio, France). F_m_ and F_m_’ were measured using saturating red pulses (3000 µmol photons m^-^² s^-1^, duration 250 ms) followed by blue light (λ = 470 nm) detection pulses. NPQ was calculated as (F_m_ - F_m_’)/F_m_’ and Φ_PSII_ as (F_m_’ - F_s_)/F_m_’ (Maxwell and Johnson, 2000).

For measurements of NPQ induction in the dark, cells were concentrated by centrifugation (5 min, 3000 g) and the pellet was resuspended in fresh ESAW at the concentration of 10^7^ cells mL^-1^. The external pH was acidified by sequential additions of acetic acid up to 4 µM, and then neutralized using potassium hydroxide up to 2 µM. Fluorescence was measured with a PAM-101 fluorimeter (Walz, Effeltrich, Germany) after pH equilibration. NPQ was calculated upon reaching steady state fluorescence levels.

### Spectroscopic measurements

Spectroscopic measurements were performed using a JTS-10 spectrophotometer (Biologics, France) equipped with a white probing LED and the appropriate interference filters (3-8 nm bandwidth).

For cytochrome *b_6_f* turnover measurements, cells were dark acclimated for 10 minutes in the dark and permeabilized with 20 µM nigericin in the dark. Acetic acid was added by steps of 1 mM to reach a pH in the external medium of about 4. Kinetics of cytochrome b_6_f reduction were measured by exposing cells to a single turnover laser flash. The cytochrome redox changes were calculated as [Cyt] = [554] - 0.4 × ([520] + [565]), where [554], [520] and [566] are the absorption difference signals at 554 nm, 520 nm and 566 nm, respectively (Bailleul et al., 2015). The turnover rate *k* was calculated as *k* = ln2/t_1/2,_ where t_1/2_ is the half time of the reduction phase.

We calibrated the pH values measured in Figure 1 by assessing the pH dependence of cytochrome (cyt) *b_6_f* turnover rate (Supplementary Figure S2). This parameter is tightly pH-controlled due to the presence of evolutionary conserved residues in the complex (Berry et al., 2000; Sarewicz et al., 2021). Its pH dependency has already been employed as a probe to deduce lumenal pH value in vivo ((Finazzi and Rappaport, 1998); see Methods). Consistent with the aforementioned hypothesis of an incomplete pH equilibration within the diatom cell, we found that the apparent pKa of the cytochrome turnover pH-dependency was shifted to more acidic pH values in *P. tricornutum* (Figure S2B, C, open circles and short dots) than in *C. reinhardtii* ((Finazzi, 2002); Supplementary Figure S2B, C). Using the cyt *b_6_f* shift (Supplementary Figure S2B, open circles) to rescale the pH (Figure 1C, upper X scale) led to a similar pH sensitivity of NPQ in *P. tricornutum* and *C. reinhardtii* (Tian et al., 2019). This pH dependency of NPQ closely resembles the pH profile of the xanthophyll de-epoxidase measured in P. tricornutum (Jakob et al., 2001).

For ΔΨ measurements, cells were acclimated to red light for 5 min in the presence of 10 µM nigericin if needed. ECS changes were measured by exposing these pre-acclimated cells to a 10 ms saturating white light pulse (1200 µmol photons m^-2^.s^-1^) and following its decay. Absorption relaxation signals ([554], [520] and [565]) were measured respectively at 554, 520 and 565 nm. The ECS signals were calculated as ECS_lin_ = 1.1 x [520] - 0.25 × [554] + 0.1 [565] and ECS_quad_ = 0.94 × [565] + 0.15 × [554] - 0.06 × [520]. ΔΨ was calculated using a parabolic equation fit as described in (Bailleul et al., 2015). ECS data were normalized to the ECS_lin_ increase upon a saturating laser flash in the presence of DCMU and HA. An increase of ΔΨ in the presence of nigericin reveals the presence of a ΔpH component of the PMF as compared to control conditions.

9-aminoacridine (9-aa) fluorescence was measured to determine ΔpH using the DUAL-ENADPH and DUAL-DNADPH units of the DUAL-PAM-100 NADPH/9-aa module (Walz Effeltrich; (Schreiber and Klughammer, 2009; Johnson and Ruban, 2011)). Excitation was provided by 365 nm LEDs and fluorescence emission was detected between 420 and 580 nm. Measurements were performed on WT and mutant *P. tricornutum* cells (5 μg chlorophyll *a* mL^-1^, at 19 °C) in the presence of 1 μM 9-aa (5-min incubation in the dark) and 50 μM diaminodurene (DAD; 2-min incubation in the dark). DAD was used to amplify the 9-aa fluorescence signal. Actinic light and saturating pulses were provided by an array of 635 nm LEDs. NPQ was induced by a 5-min of illumination with a 472 μmol photons m^-2^ s^-1^ actinic light, and recovery was monitored in the dark. The extent of 9-aa fluorescence quenching was assessed after correction of the 9-aa signal drift observed. Measurements were repeated on 3-4 biological replicates per genotype.

### Pigment analysis by HPLC

Cells were harvested during mid-exponential phase at a concentration around 5 million cells/mL. They were placed on plates and exposed to light until steady state NPQ was reached. Cells were harvested, immediately frozen in liquid nitrogen and dehydrated overnight. The resulting powder was resuspended in 1.2 mL 100% methanol in the dark, and always kept at 4°C and under argon atmosphere. After centrifugation at 4000 g for 2 min, the supernatant was dried under argon flux. The remaining pigments were solubilized in 100 µL N,N-dimethylformamide (DMF) and analyzed by HPLC on a C18 column (Macherey-Nagel, France) as indicated in (Allorent et al., 2013). The following solvents were used: (1) methanol 80% v/v ammonium acetate 100 mM (2) 90% acetonitrile v/v (3) ethyl acetate. Pigments were identified according to their retention time and absorption spectrum.

The de-epoxidation rate (D.E.S.) was calculated as 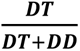 (DT = diatoxanthin; DD = diadinoxanthin. A linear regression provided the following relationship between NPQ and D.E.S.: NPQ = 14.158 × D.E.S (R² = 0.93).

### Mathematical model

A kinetic model of NPQ was built to test various hypotheses regarding the proton regulatory circuit in diatoms. This model was adapted to diatoms from a pre-existing kinetic model of NPQ designed to study the effect of short-term light memory in *Arabidopsis thaliana* (Matuszyńska et al., 2016). A general scheme of the model of the electron transport chain is shown in Figure 3A. Some important changes were made to consider physiological differences between diatoms and plants and are summarized below.

The model comprises a set of five ordinary differential equations (ODEs) following the dynamics of the reduced state of the plastoquinones (PQ), the stromal concentration of ATP, the lumenal proton concentration (H), the fraction of epoxidized xanthophylls (DD), and the fraction of active ATPase. Lumen and stroma volumes in diatoms are of a similar range (Uwizeye et al., 2021), in contrast to plants, where stromal volume is several-fold bigger. To account for a slower effect of proton accumulation, we took the simple approach of simply changing the pH function calculating the lumenal pH to account for a similar volume as the one of the stroma.

In this model, we adopted the same stoichiometries to characterize the activity of PQ, lumenal H, ATP and ATPase as introduced by Matuszyńska (Matuszyńska et al., 2016). Photosystem II was described as in the original model linking together chlorophyll excitation with the occupation of reaction centers (RC), resulting in a four-state description. The four states were determined by a quasi-steady-state approximation and solved numerically during the integration.

To test the hypothesis that PtKEA3 is involved in the regulation of the ΔpH by exchanging K^+^/H^+^ ions across the thylakoid membranes, we introduced the partitioning of the Proton Motive Force (PMF) between its ΔΨ and ΔpH components. To model the rate of PtKEA3 activity with the rate constant proportional to the ratio of ΔpH and ΔΨ, we introduced a new constant (k_KEA3_) accounting for the PMF partition between ΔpH and ΔΨ and a new derived variable (H_β_). While protons generated by linear electron flow (H^+^) are both osmotic and charged species (acting both on ΔpH and ΔΨ), the variable H_β_ encapsulates only the osmotic component of protons. Consequently:

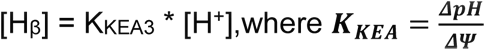

The rate of ΔpH to ΔΨ conversion was hence defined as v_KEA3_ = k_leak_ * ΔpH(H_β_ ) In plants, a four-state quenching model had been adopted to account for the uncoupling between PSBS activation, violaxanthin formation and NPQ. The data presented here and in the literature (Goss et al., 2006; Blommaert et al., 2021) on the other hand suggest that diatoxanthin formation is always concomitant with NPQ. To account for this difference between plants and diatoms, we hypothesized that the quencher (Q) formation depends linearly on the de-epoxidation state of the xanthophyll cycle, with Q calculated as:

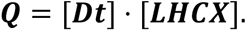

Moreover, the regulation of the xanthophyll de-epoxidase activity depends on changes in the ΔΨ and ΔpH ratio and is described by the equation below:

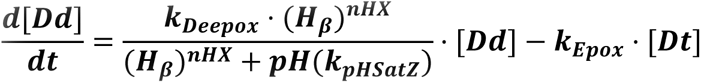

With the assumption that the rate of de-epoxidase k_Deepox_ is two times faster than the epoxidase k_Epox_. Since the homologues of these enzymes are known to respond to pH changes rather than electrical changes, the variable H_β_ was considered in this equation.

The version of the model described in this manuscript has been implemented in Python using modelbase v1.2.3 software (van Aalst et al., 2021) and integrated using provided assimulo solvers. Parameter values used for the simulation are provided in Supplementary Table S3..

Fluorescence was calculated as the probability that the excited state of chlorophyll reverts back to its ground state emitting fluorescence. Fluorescence can be emitted from two of the four states of photosystem II (B_0_ and B_2_) and the signal is also proportional to the occupation of these two ground states. Overall, the fluorescence is derived using the formula:

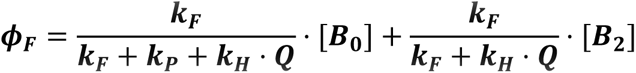

where k_F_, k_H_ and k_P_ are the rates of fluorescence emission, quenching (k_H_) and photochemistry (k_P_), respectively. The parameters were adjusted so the photosynthetic efficiency 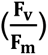 in a dark-adapted state ranges between 0.6 – 0.7, corresponding to the commonly observed values in diatoms (Taddei et al., 2016):

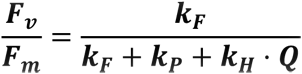

NPQ is calculated from fluorescence using the formula:

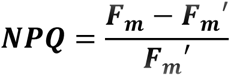

Where *F_m_* is the maximum fluorescence emission levels in the dark and *F_m_*′ is the maximum fluorescence during actinic light exposure, measured by a saturating pulse of light.

The code is published under open license in the GitLab repository: https://gitlab.com/qtb-hhu/models/npq-diatoms2020.

## Accession numbers

Supplementary Table S5

## Author Contributions

C.S., M.S., V.G., H.H., B.B., F.C. G.A. and G.F. designed research, C.S., M.S., V.G. A.M., E.G. X.Z. C.G., Y.P., J.A., H.H., F.C., G.A. and G.F. performed research; C.S., M.S., V.G., A.M., X.Z. C.G., H.H. A.V.R., B.B., F.C. G.A. and G.F. analyzed data; C.S., V.G., A.M., and G.F. wrote the manuscript.

## Acknowledgments

The authors would like to thank Dr Leonardo Magneschi (Ingenza Limited, UK), and Prof Oliver Ebenhöh (University of Dusseldorf, De) for discussions and help in the first steps of this project. We would like to thank Dr Marcel Kunz (CEA Grenoble) for technical help with the HPLC facilities. This project received funding from the LabEx GRAL (ANR-10-LABX-49-01) and was financed within the University Grenoble Alpes graduate school (Ecoles Universitaires de Recherche; CBH-EUR-GS) (ANR-17-EURE-0003). G.F., M.S. and C.G. acknowledge the support by the European Research Council (ERC) Chloro-mito (Grant No. 833184). E.G. and G.F received funding from the French Research Funding agency (ANR) grant ‘Momix’ (Projet-ANR-17-CE05-0029). G.A. and G.F received funds from the HFSP ‘Photosynthesis light 37tilization dynamics and ion fluxes: making the link’. X.Z and G.F. received funds form a CEA grant on carbon circular economy. G.A. received funds from the CNRS ‘Momentum’ program. V.G. received funds from The Leverhulme Trust (RPG-2018-199). A.V.R. received funds from The Leverhulme Trust (RPG-2018-199), the Biotechnology and Biological Sciences Research Council (BB/R015694/1) and The Royal Society Wolfson Research Merit Award (WM140084). A.M. received funding from the Deutsche Forschungsgemeinschaft (DFG, German Research Foundation) under Germany’s Excellence Strategy – EXC-2048/1 – project ID 390686111 and DFG Research Grant MA 8103/1-1. B.B. acknowledges the support by the European Research Council (ERC) PhotoPHYTOMIX project (Grant No. 715579).

## Supplementary Information for

**Supplementary Figure S1.**
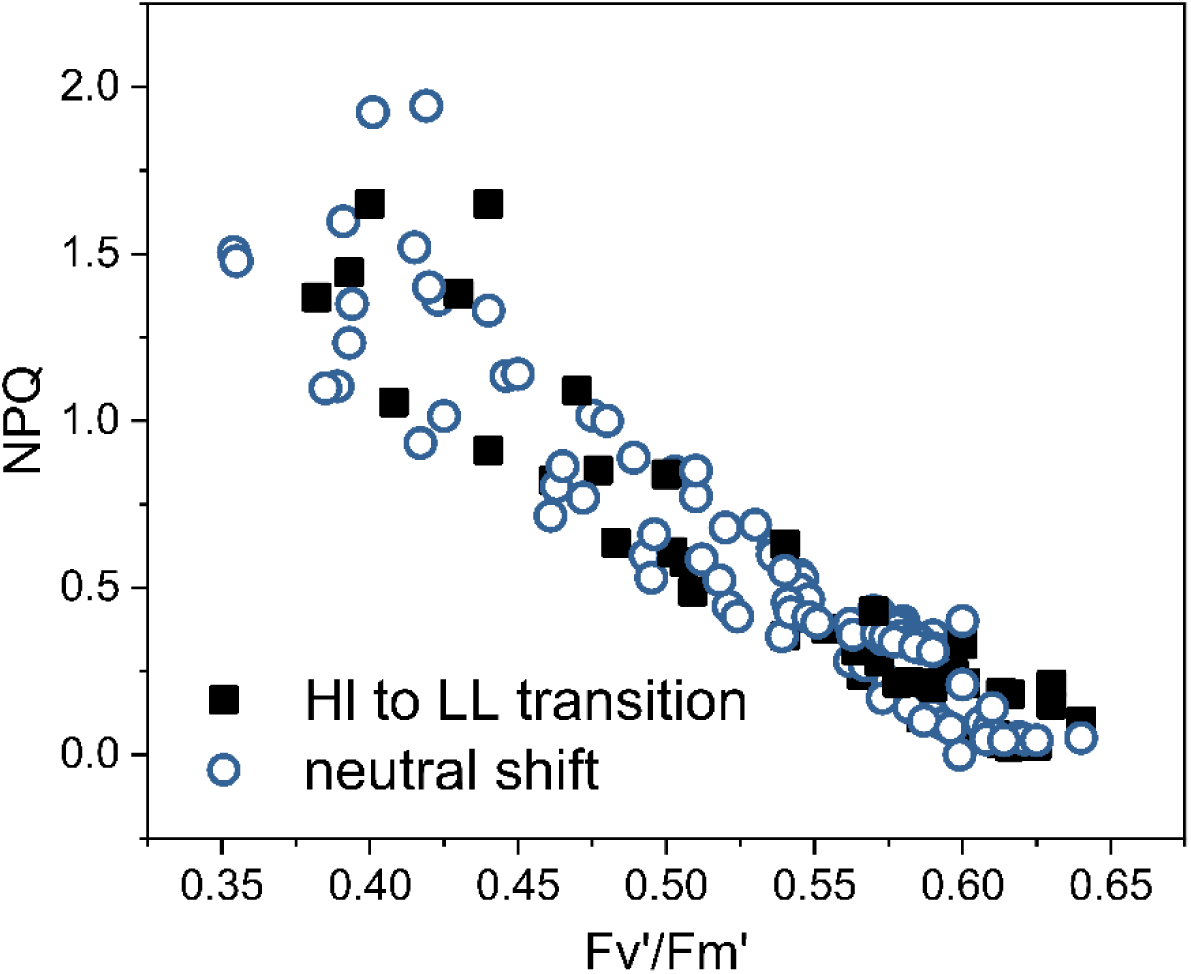
pH Induced fluorescence quenching does not affect the Stern-Volmer relationship between NPQ and PSII photochemical quantum yield (supports Figure 1). The PSII photochemical capacity was evaluated via the fluorescence parameter Fv’/Fm’ (Maxwell and Johnson, 2000) during the HL to LL relaxation of NPQ and at different pH during both neutralization (blue circles) of the medium pH. N= 4

**Supplementary Figure S2.**
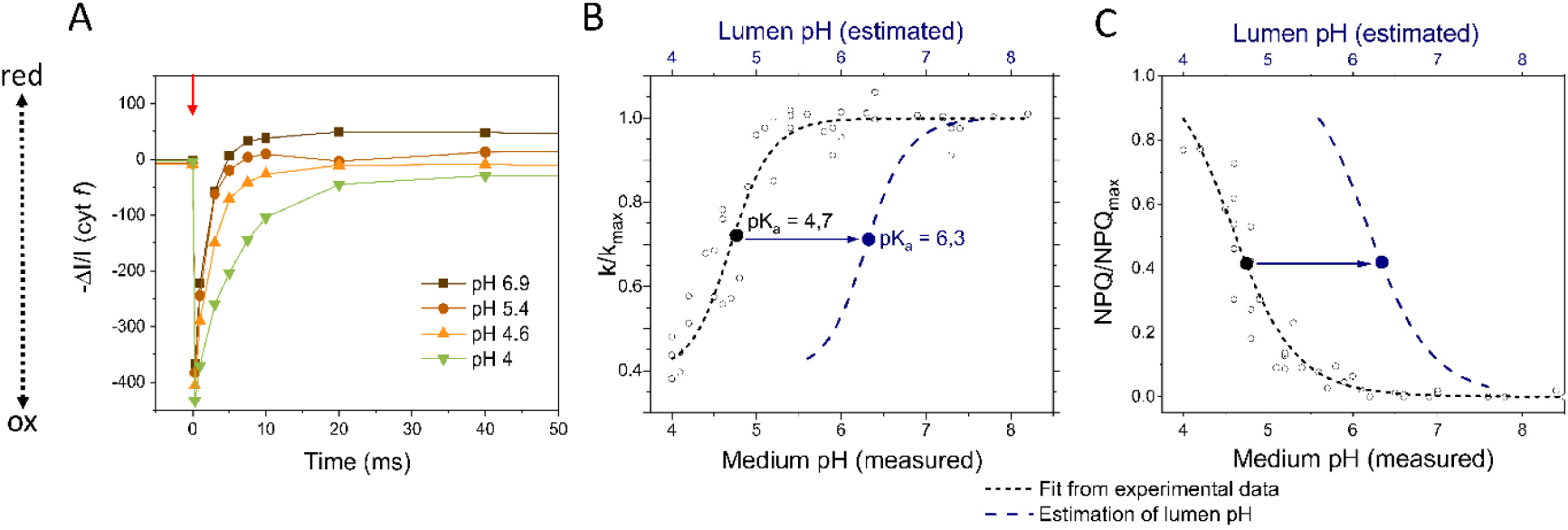
Calibration of the lumenal pH in *P. tricornutum* cells as a function of medium acidification (supports Figure 1). (**A**) Rate of cytochromes c_6_ + *f* reduction after a saturating turnover flash (red arrow) at different pH in the presence of 20 µM nigericin. (**B**) The ratio between the rate constant measured at every pH and the maximum one (k/k_max_, open circles) was fitted with a sigmoidal equation to identify the apparent pKa of 4,7 (black dotted line). This parameter is tightly pH-controlled due to the presence of evolutionary conserved residues in the respiratory *bc*_1_ and photosynthetic *b_6_f* complexes (Berry et al., 2000; Sarewicz et al., 2021). Thus, by comparing it with the previously published pKa calculated in *C. reinhardtii* (6.3 (Finazzi, 2002), we estimated the shift between the measured pH (lower X axis) and the luminal pH attained during the titration (upper X axis, blue dashed line). Thanks to this parameter we inferred the possible shift in the pH dependence of NPQ as 6,3 - 4,7 = 1,6 pH units (**C**). pH equilibration was performed as described in Fig. 1.

**Supplementary Figure S3.**
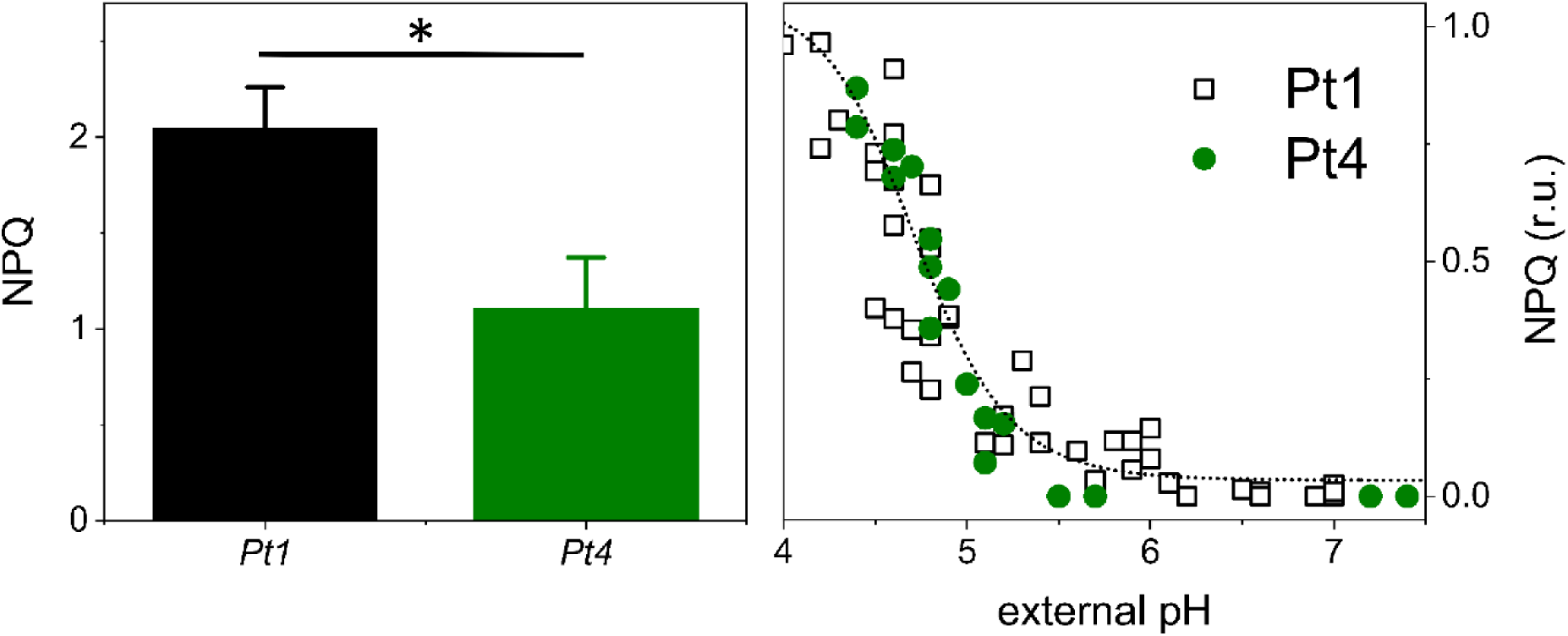
NPQ extent (left) and pH dependency (right) in the *P. tricornutum* Pt1 and Pt4 ecotypes (supports Figure 1). Same experimental design as shown in Fig. 1. N = 3 mean ± SD. In the right panel, NPQ amplitudes in Pt1 and Pt4 have been normalized to facilitate their comparison. Data represent traces from 3 biological replicates. Asterisks indicate significant differences in NPQ between the Pt1 and Pt4 ecotypes (P < 0.05).

**Supplementary Figure S4.**
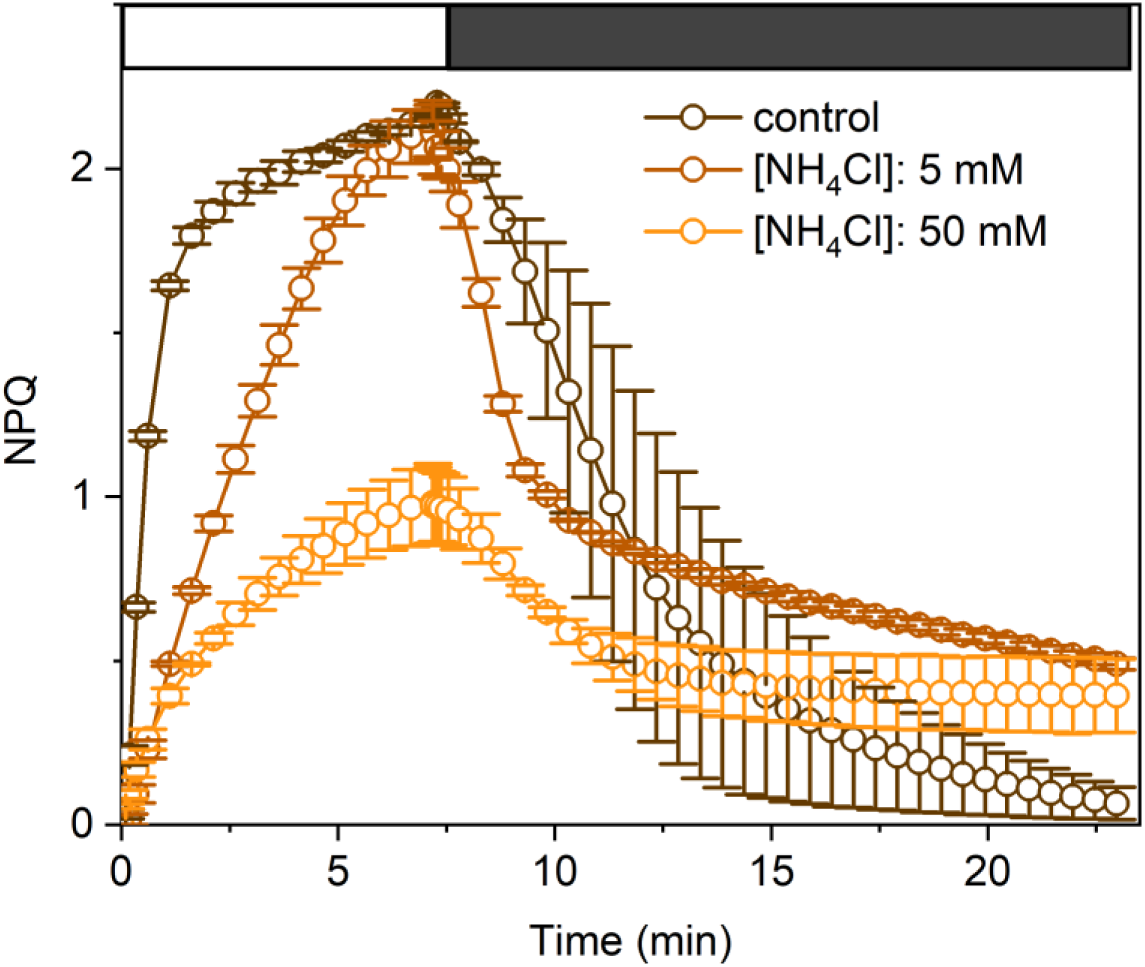
NH_4_Cl sensitivity of NPQ in *P. tricornutum* cells (supports Figure 1). Different concentrations of NH_4_Cl were added before measuring NPQ induction in HL (1200 µmol photons m^-2^ s^-1^, white box) and relaxation in LL (25 µmol photons m^-2^ s^-1^, grey box). N = 3 mean ± SD.

**Supplementary Figure S5.**
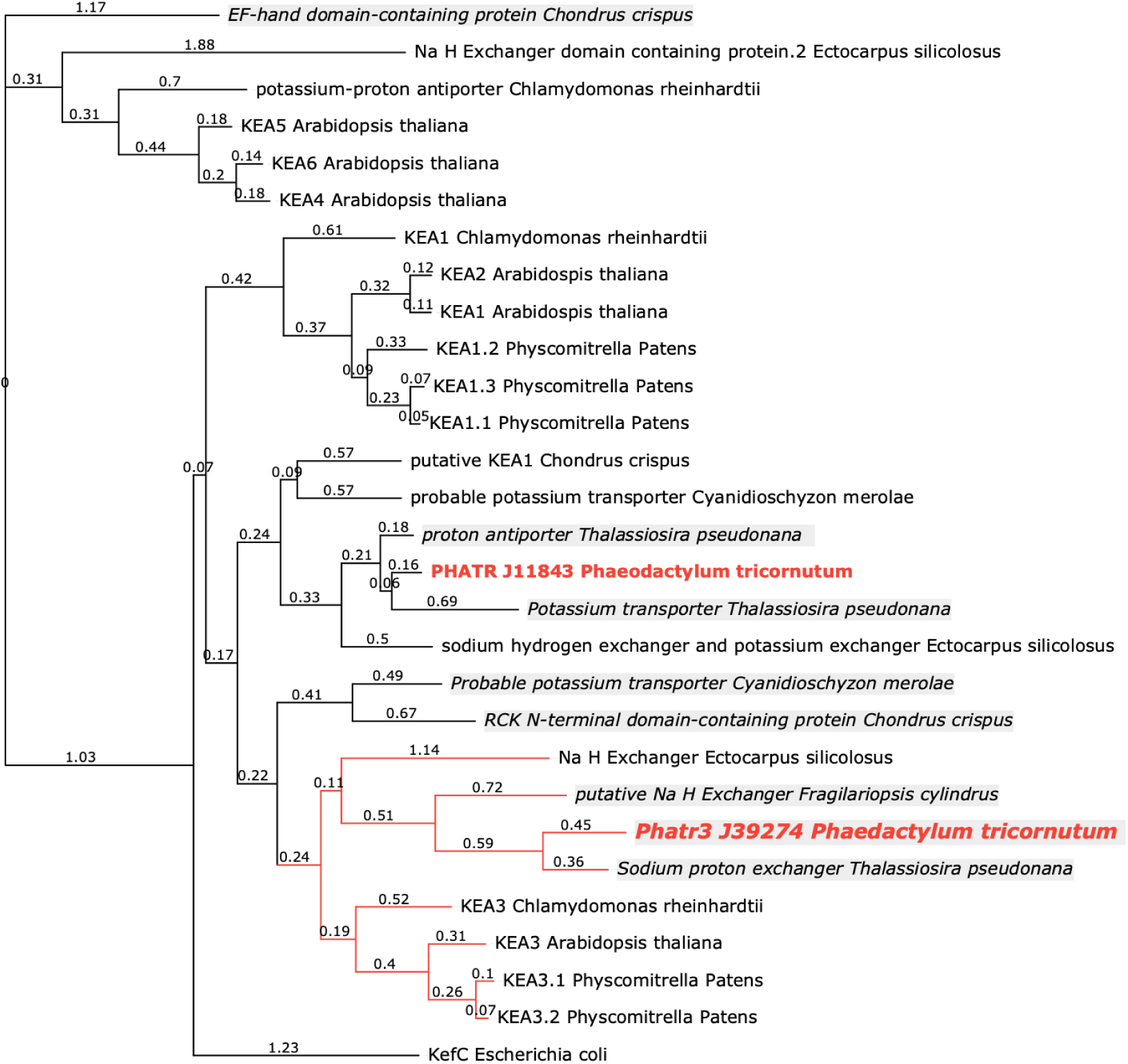
KEA mid-point rooted phylogeny inferred using maximum likelihood analysis (supports Figure 2). Maximum clade credibility trees summarized using TREE (https://www.genome.jp/tools-bin/ete) with the scale bar indicating substitutions per site. Branch supports: Chi2-based parametric values returned by the approximate likelihood ratio test. Two putative gene products belonging to the KEA family in *P. tricornutum*, i.e. Phatr3_J11843 and Phatr3_J39274, are marked in red. Branches in red represent putative gene products of KEA3 orthologues. Based on similarity with the *A. thaliana* KEA3 gene, Phatr3_J39274 was identified as the most likely homologue of KEA3 in *P. tricornutum.* Gene products containing a predicted EF-hand are marked in italic and highlighted in grey boxes.

**Supplementary Figure S6.**
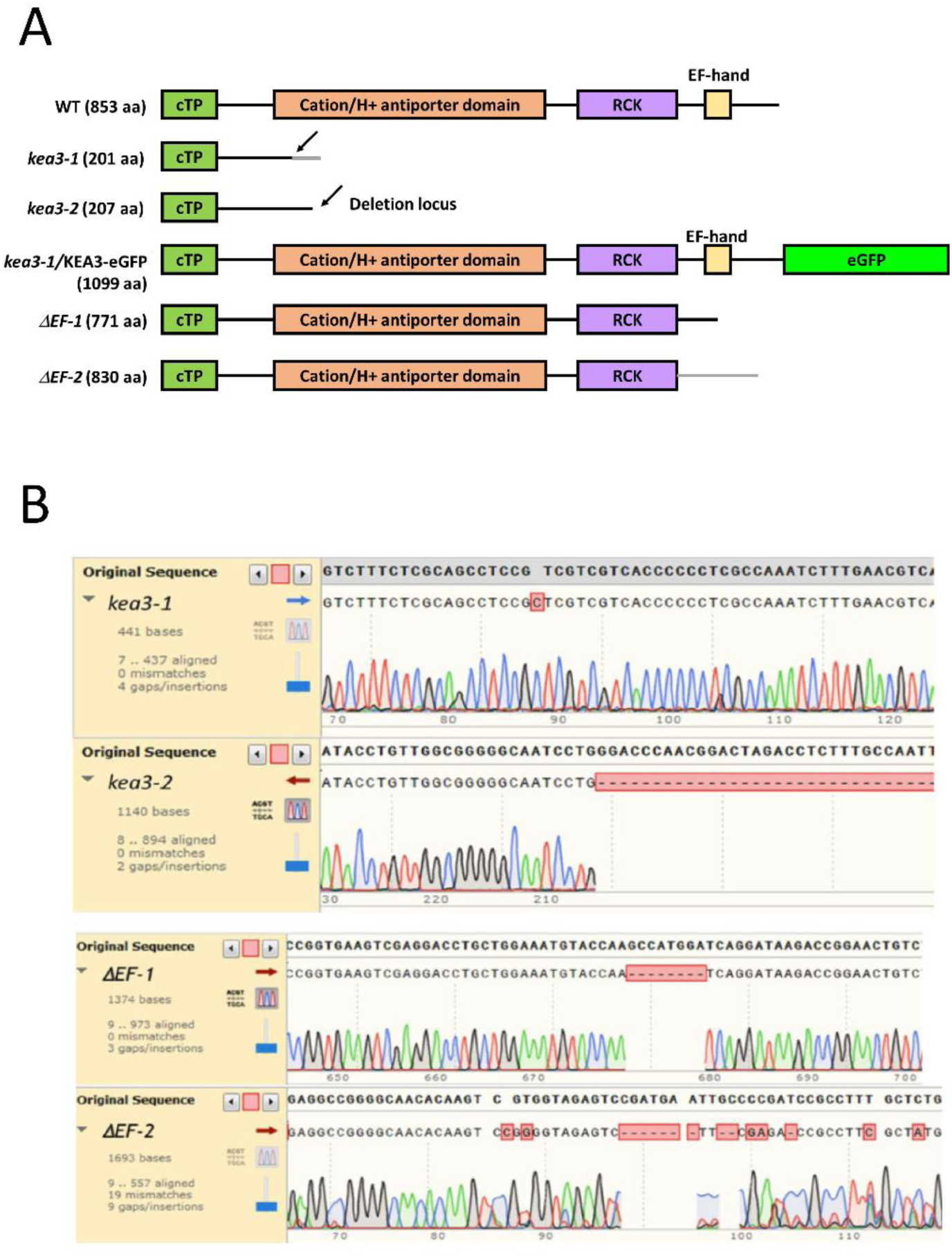
Molecular characteristics of the KO mutant genotypes employed in this work (supports Figure 2). (**A**): Schematic representation of the different domains of the KEA3 gene in *P. tricornutum*: the plastid transit peptide (green), the putative cation/H^+^ antiport (orange), the regulatory of K+ conductance (purple) and the EF-hand Ca^2+^ binding motif (yellow). Their presence/modification in the different genotypes characterized in this work is shown. Arrow heads indicate the localization of the stop codons in the nucleotide sequence of the KO mutants. (**B**) Sequencing of mutant clones. An insertion at position 484 *(kea3-1)* and a 457 bp deletion at position 558 *(kea3-2)* from the translation initiation base lead to truncated proteins of 201 *(Ptkea3-1)* and 191 *(Ptkea3-2)* aminoacids. The *ΔEF-1* and *ΔEF-2* mutations generate deletions of 8 and 23 aa, affecting the EF domain of the protein.

**Supplementary Figure S7.**
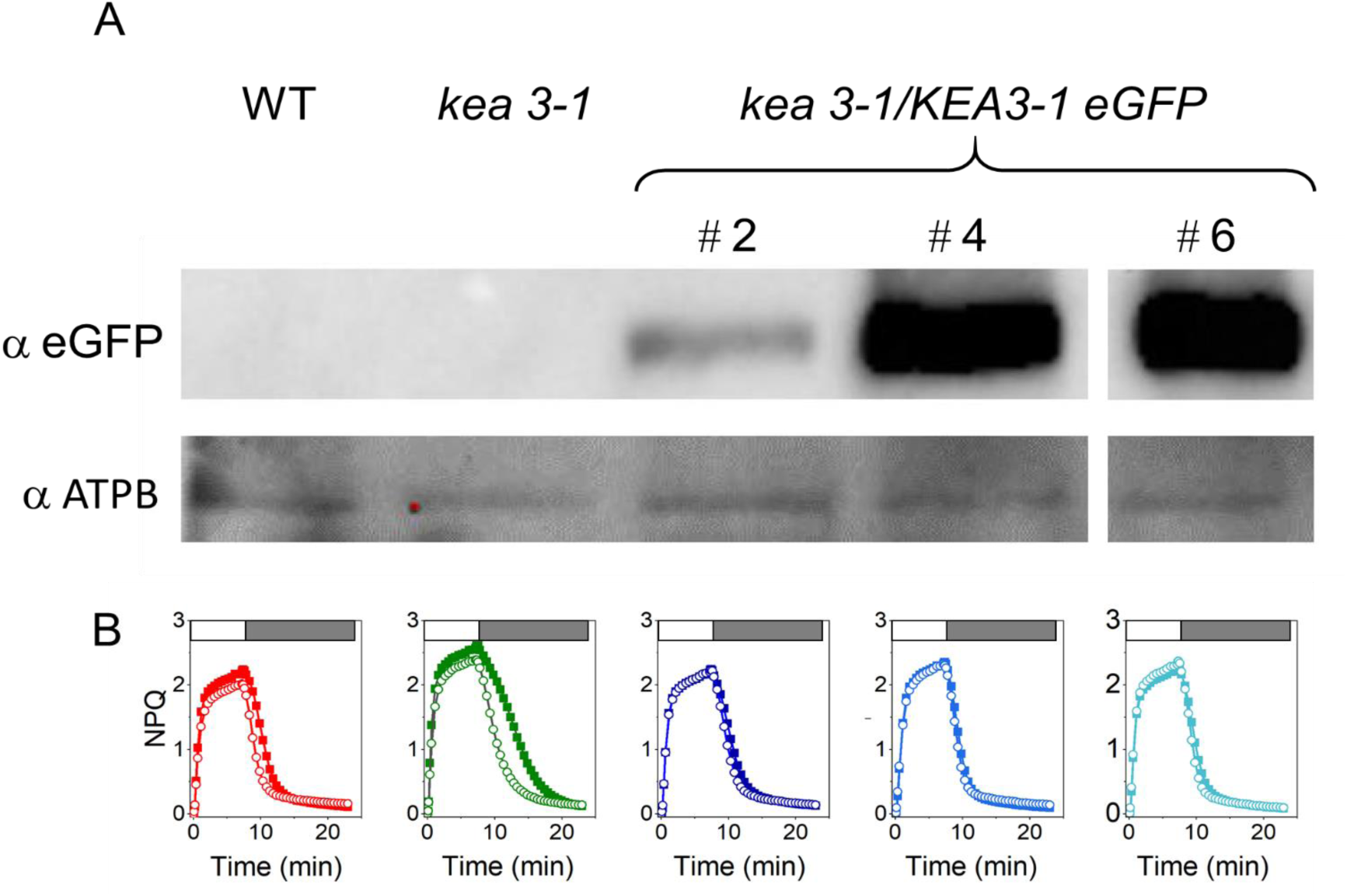
Molecular and physiological characterization of the complemented *kea3-1*/KEA3-eGFP genotype (supports Figure 2). Immunodetection analysis of KEA3 polypeptide in the WT, the *kea3-1* mutant and in 3 clones where KEA3 function was recovered upon complementation of the *kea3-1* mutant with a plasmid containing the native gene fused to the eGFP coding sequence. The #4 clone, which over-accumulates KEA3, was used in this study. The ß subunit of the ATP synthase (ATPB) was used as loading control. (B) NPQ phenotypes of the WT (red) *kea3-1* (green) and complemented *kea3-1/KEA3-*eGFP genotype (blue) in the absence (solid symbols) and presence (open symbols) of nigericin 10 µM.

**Supplementary Figure S8.**
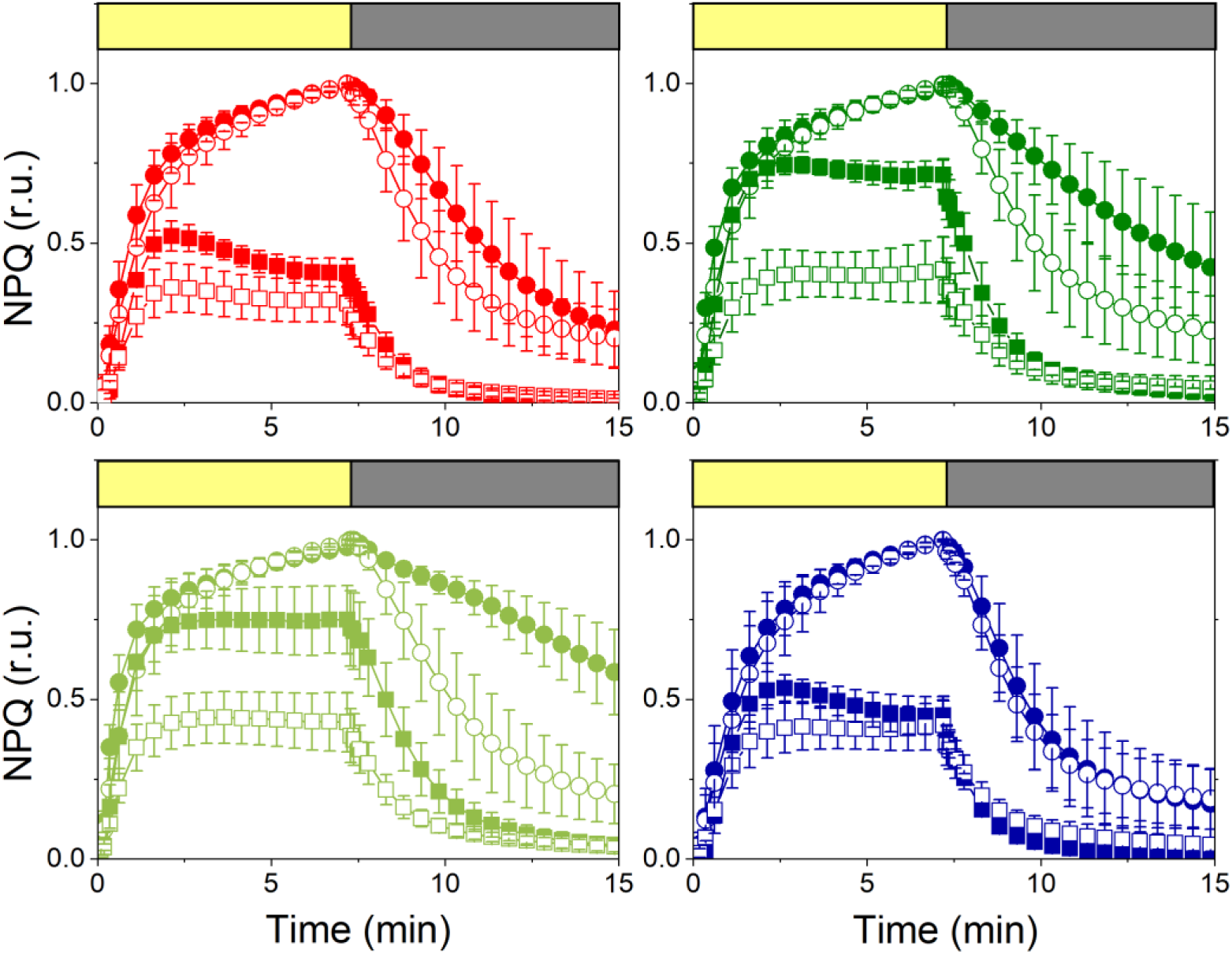
NPQ features of the *kea3-1* and *kea3-2* KO mutants compared to the WT and kea3-1/KEA3-eGFP genotypes (supports Figure 2). NPQ features in *kea3-1* (dark green) and *kea3-2* (light green) compared to the WT (red) and the kea3-1/KEA3-eGFP (blue) genotypes under moderate light (ML: 125 µmol photons m^-2^ s^-1^, squares) and high light (HL: 450 µmol photons m^-2^ s^-1^, circles) in the absence (solid symbols) or the presence (open symbols) of nigericin (10 µM). Same conditions as in Fig. 2. NPQ data were normalized to the value measured in HL. Yellow box: light on; grey box: light off.

**Supplementary Figure S9.**
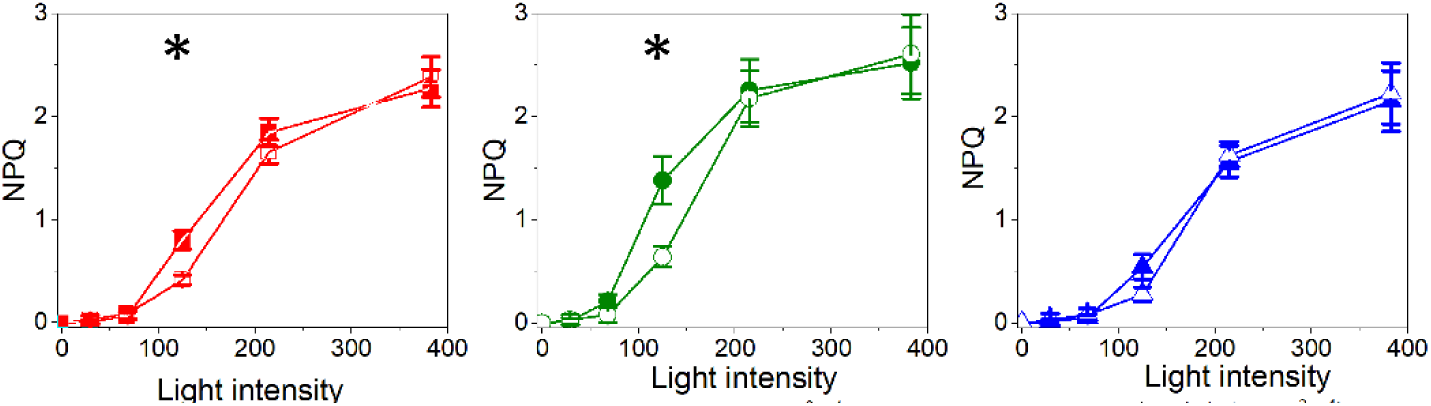
Light dependency of NPQ in WT (red) *kea3-1* and *kea3-2* KO (green) and *kea3-1/KEA3-*eGFP OE (blue) genotypes (supports Figure 2). NPQ was assessed from steady state chlorophyll fluorescence signals measured at different light intensities. Data from the two KO mutants (*kea3-1* and *kea3-2*) were averaged. Solid symbols: control; open symbols: nigericin (10 µM). N = 3 mean ± SD. Asterisks indicate significant differences between control and nigericin treated NPQ values (P < 0.05).

**Supplementary Figure S10.**
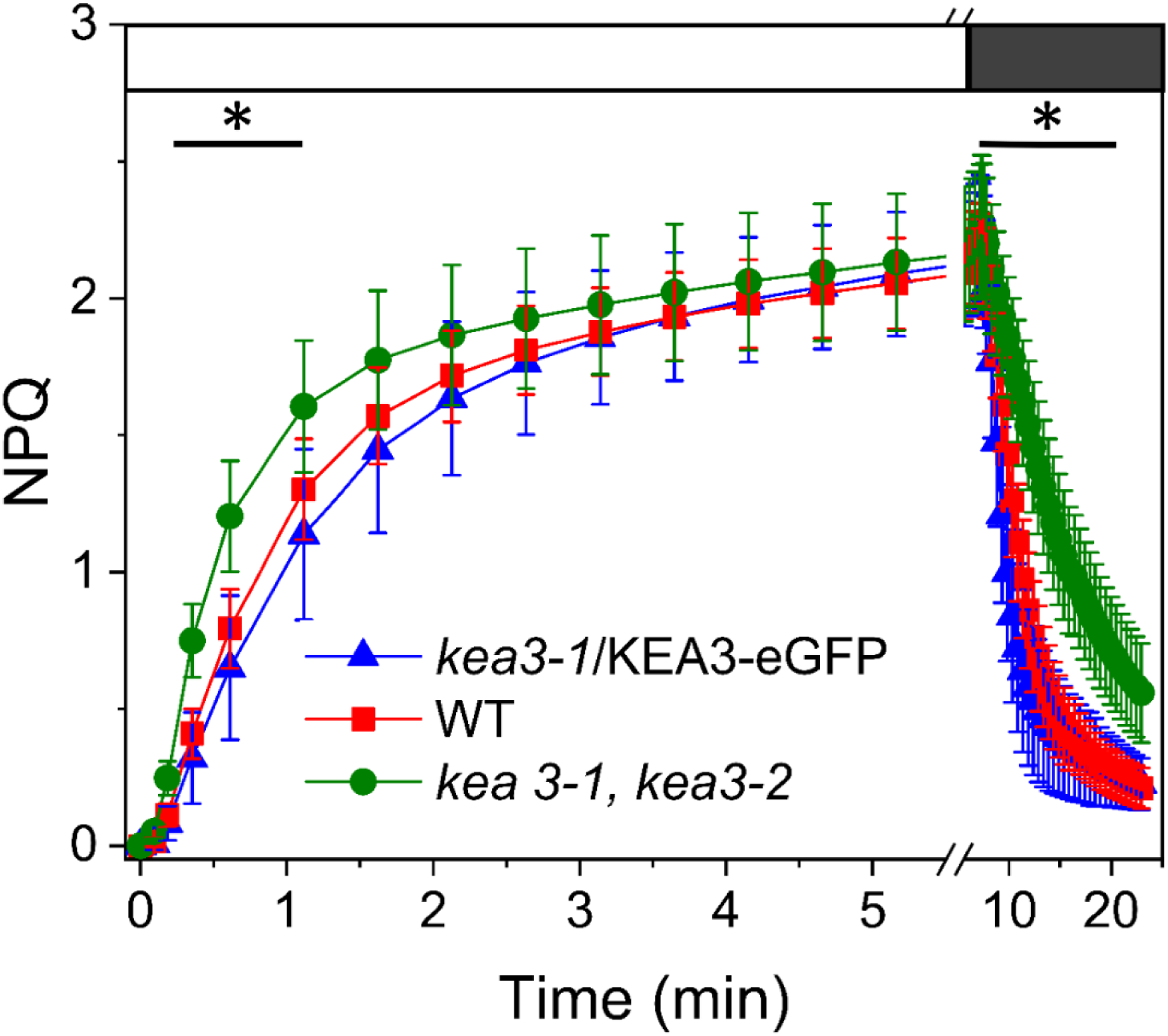
NPQ transients in WT (red), *kea3-1* and *kea3-2* KO mutants (green) and *kea3-1/KEA3-*eGFP (blue) genotypes (supports Figure 2). Same NPQ kinetics in response to HL (1200 µmols photons m^-2^ s^-1^) as in Fig. 2, presented on a different time scale. N = 15 mean ± SD. Asterisks indicate significant differences in NPQ kinetics between the *kea3-1* and *kea3-2* KO mutants, relative to the WT and the *kea3-1/KEA3-*eGFP OE lines (P < 0.05).

**Supplementary Figure S11.**
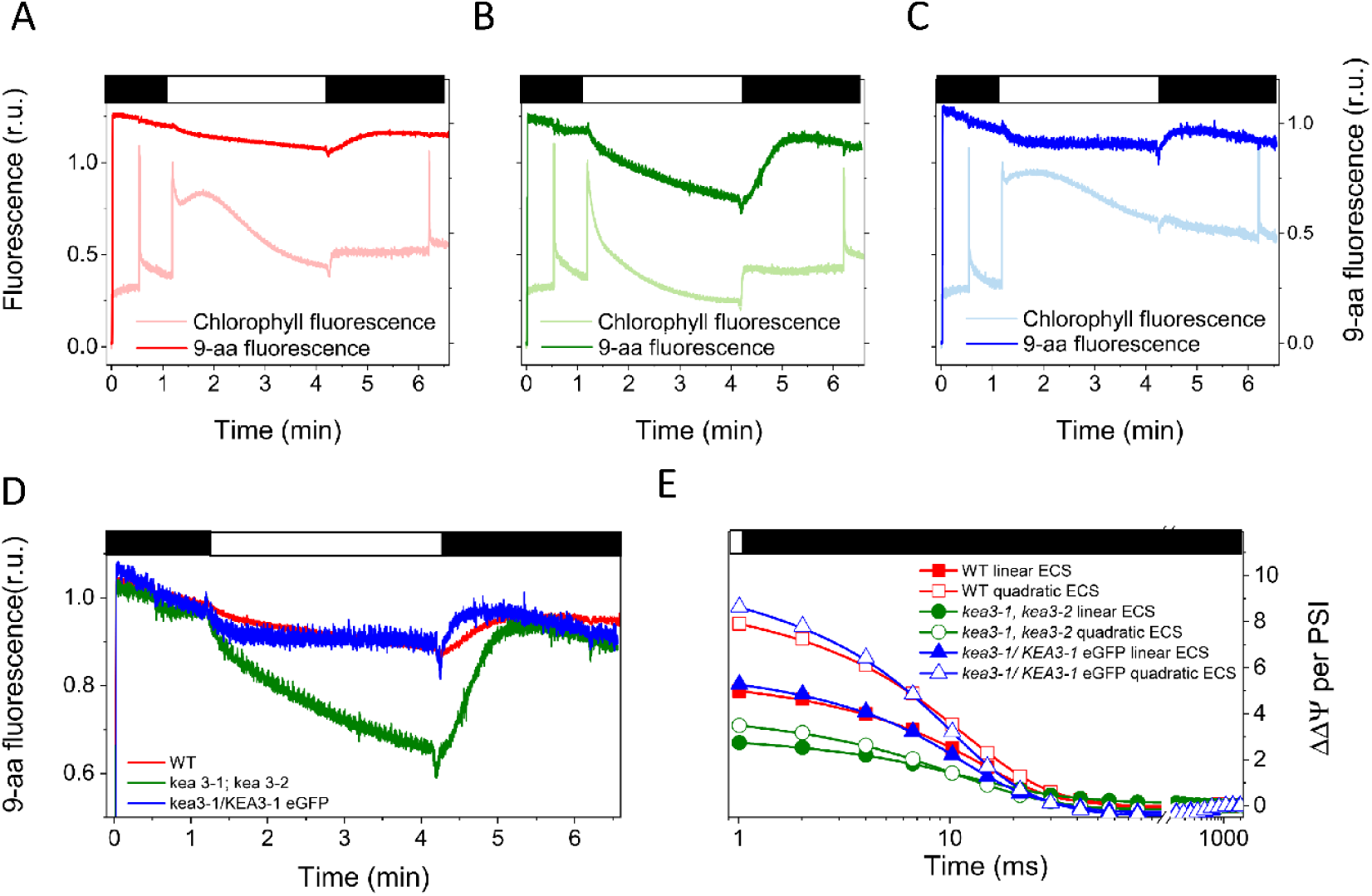
Characterization of the Proton Motive Force (PMF) in living cells of *P. tricornutum* (supports Figure 2). Chlorophyll fluorescence transient and 9-aminoacridine (9-aa) fluorescence quenching in WT **(A)**, *kea3-1* and *kea3-2* KO mutants **(B)** and kea3-1/KEA3-eGFP **(C)** cells. Chlorophyll fluorescence and 9-aa quenching were measured with a DUAL-PAM-100. White box: actinic light (472 µmol photons m^-2^ s^-1^) on. Black box: actinic light off. Representative chlorophyll fluorescence and 9-aa traces from 3-4 biological replicates with similar results. (**D**) Comparison of ΔpH estimates assessed by 9-aa quenching measurements, between WT (red), KO (green) and OE (blue) genotypes. (**E**) Measurements of the ΔΨ using the Electro Chromic Shift (ECS). Traces represent the light to dark relaxation of the linear (solid symbols) and quadratic (open symbols) ECS in WT (red), KO (green) and OE (blue) genotypes. Representative traces of an experiment repeated 3 times with similar results. Actinic light was 590 µmol photons m^-2^ s^-1^.

**Supplementary Table S1.**
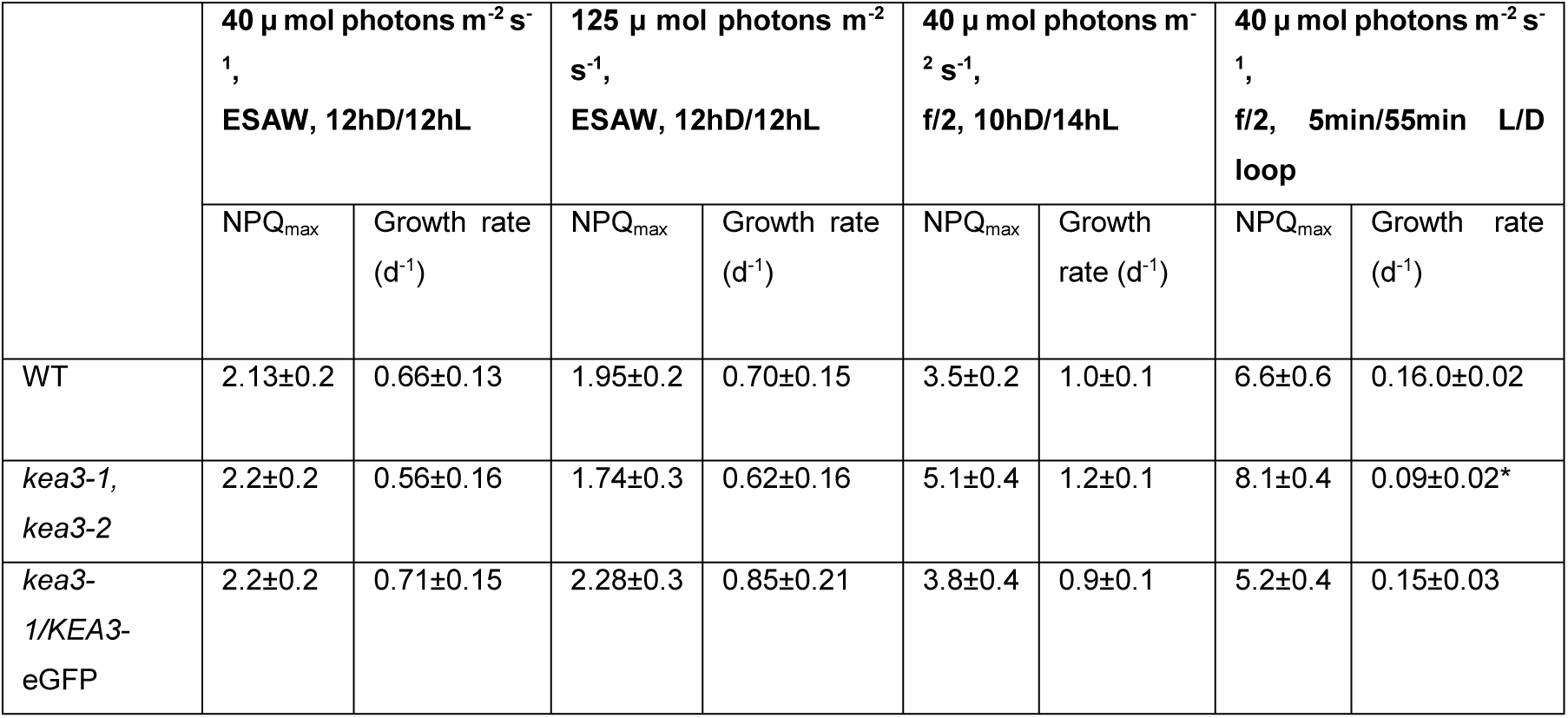
NPQ features and growth rates of *P. tricornutum* cells grown under different light and nutrient conditions (supports Figure 4). Cells grown in the f/2 medium turned out to be more sensitive to inhibition by protonophores. N= 3 mean ± SD. The asterisk indicates significant differences in growth rate between the *kea3-1* and *kea3-2* KO mutants relative to the WT and *kea3-1/KEA3-*eGFP OE lines (P < 0.05).

**Supplementary Table S2.** Protonophore sensitivity of NPQ in WT cells grown in ESAW and f/2 media (supports Figure 4). N = 5, Mean ± SD.

| 40 $\mu$ mol photons $\text{m}^{-2} \text{s}^{-1}$,<br>ESAW, 12hD/12hL | | 40 $\mu$ mol photons $\text{m}^{-2} \text{s}^{-1}$<br>f/2, 10hD/14hL | |
| --- | --- | --- | --- |
| NPQ decrease<br>(5 mM $\text{NH}_4\text{Cl}$) | NPQ decrease<br>(100 $\mu\text{M}$ Nig.) | NPQ decrease<br>(10 mM $\text{NH}_4\text{Cl}$) | NPQ decrease<br>(100 $\mu\text{M}$ Nig.) |
| 25% ($\pm 15\%$) | 50% ($\pm 10\%$) | 91% ($\pm 1.3$) | 94% ($\pm 2.9$) |

**Supplementary Table S3:**
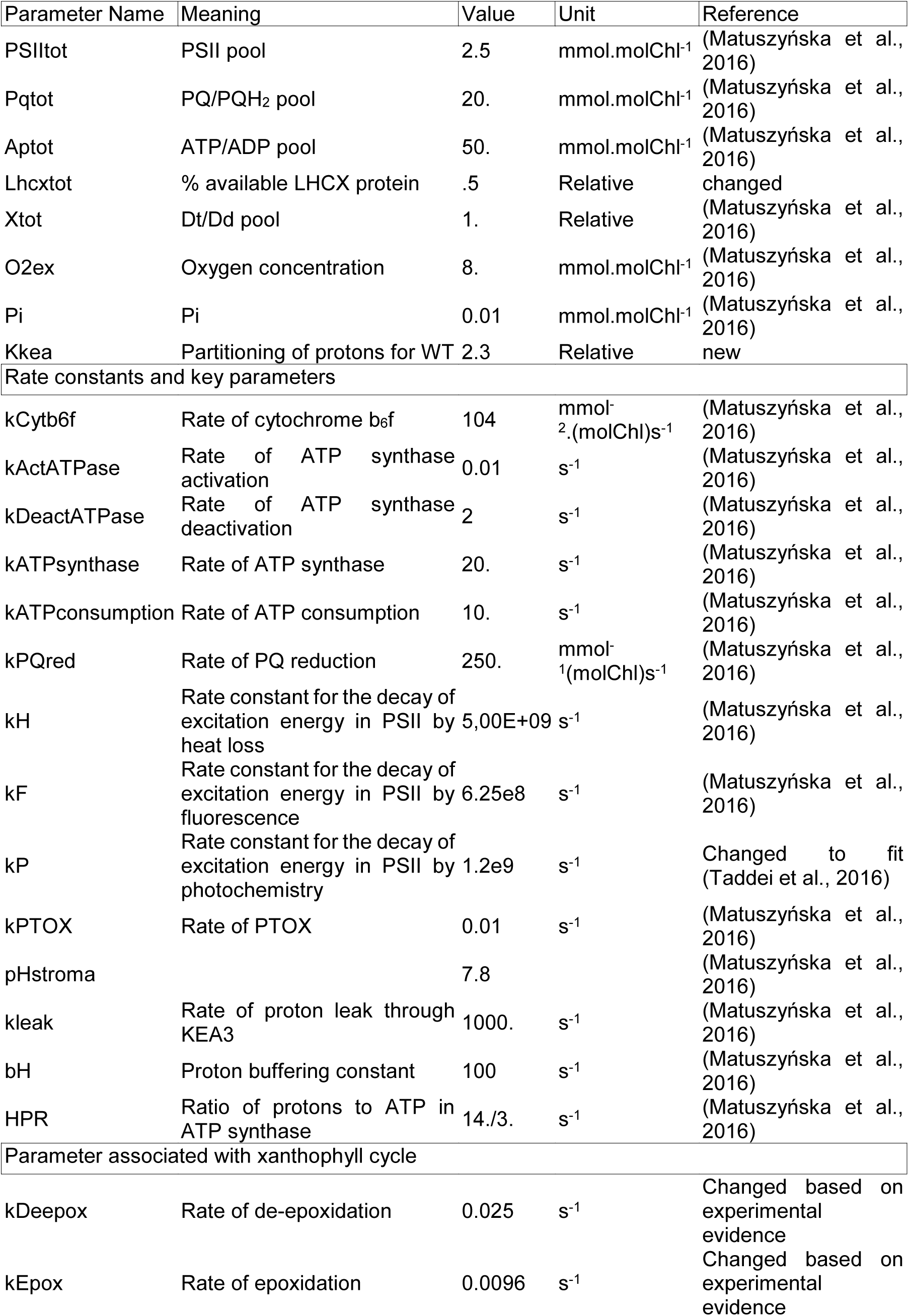
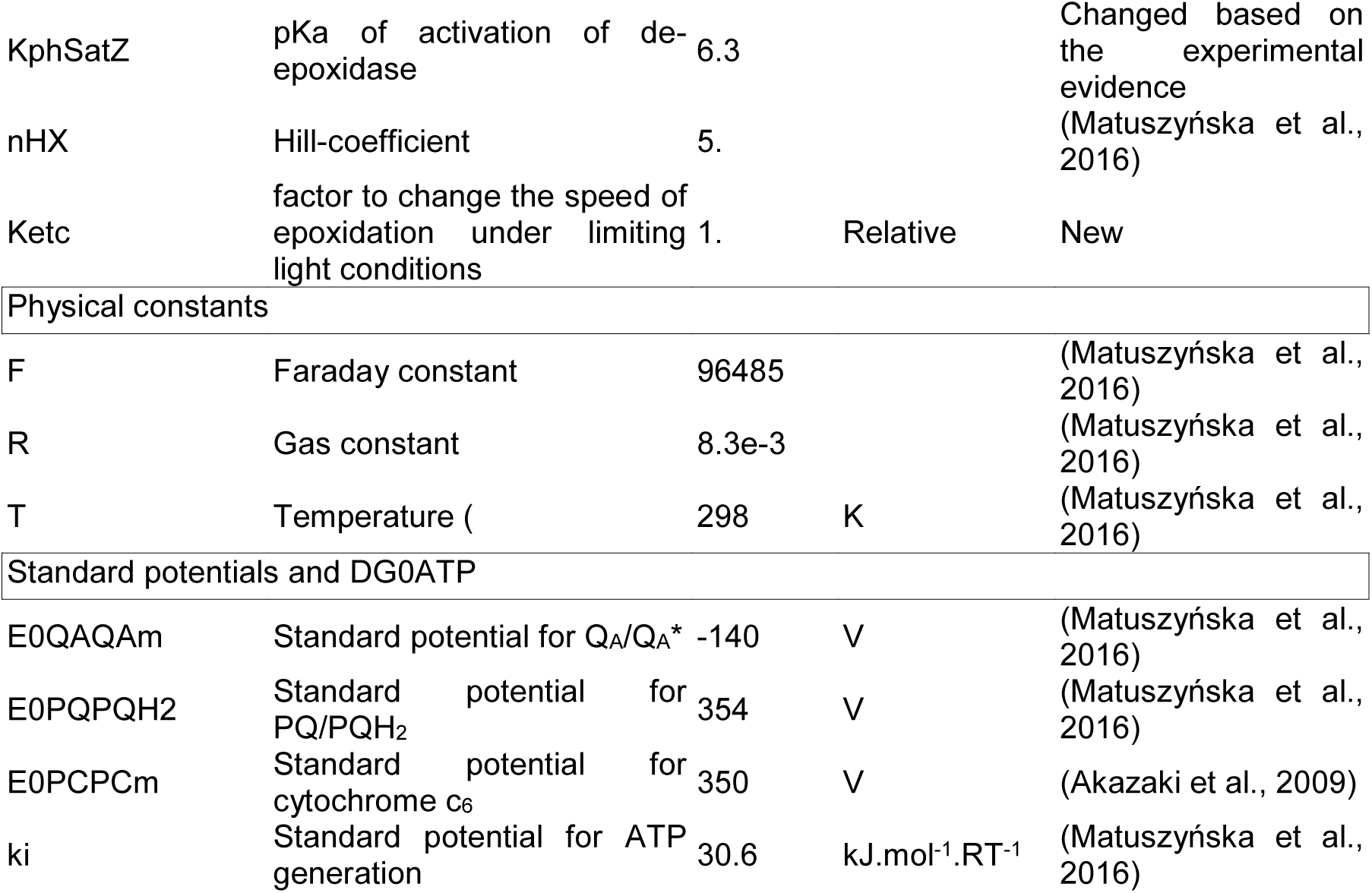
Parameters used in the model simulations (Supports Figure 3).

**Supplementary Table S4:**
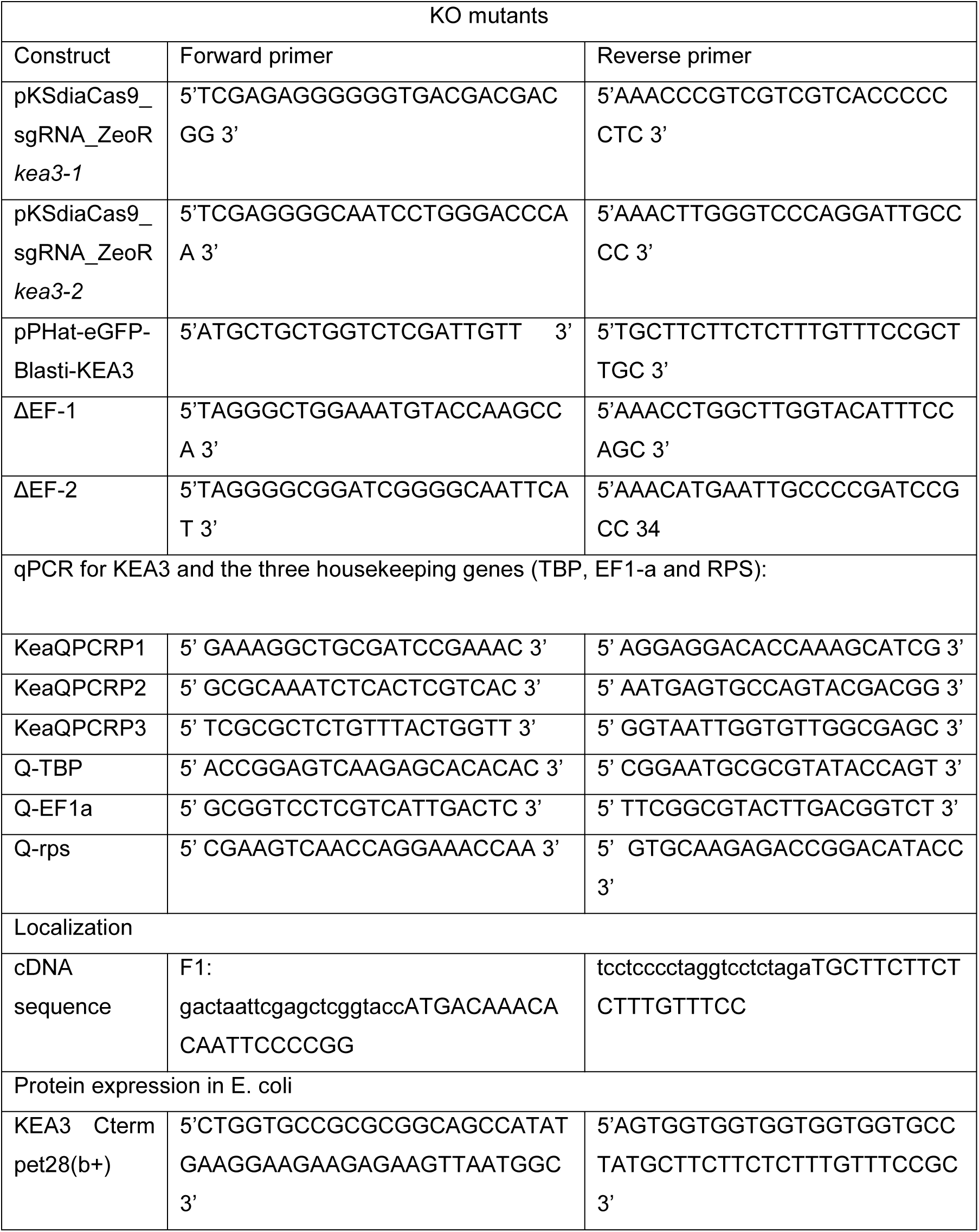
List of primers used in this study.

**Supplementary Table S5:**
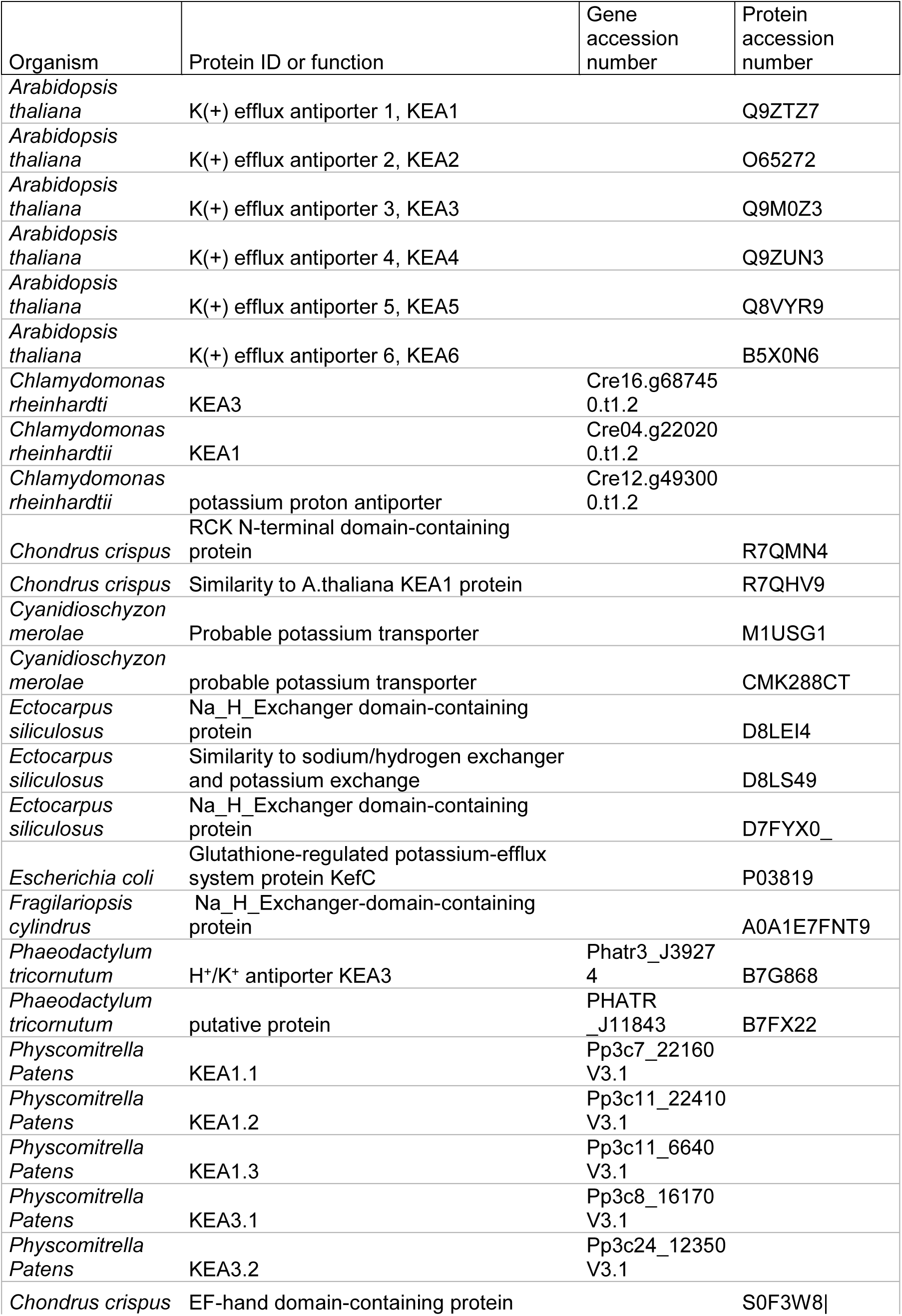
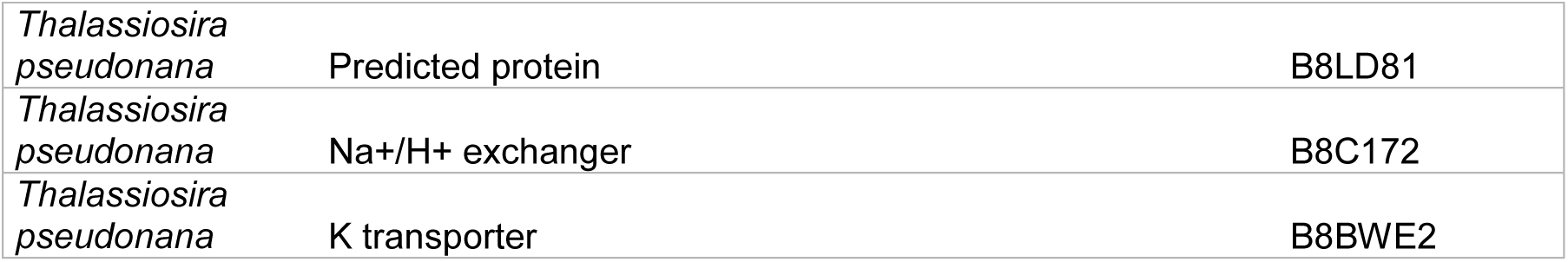
Accession numbers of proteins used in this study.

